# AI-guided discovery of the invariant host response to viral pandemics

**DOI:** 10.1101/2020.09.21.305698

**Authors:** Debashis Sahoo, Gajanan D. Katkar, Soni Khandelwal, Mahdi Behroozikhah, Amanraj Claire, Vanessa Castillo, Courtney Tindle, MacKenzie Fuller, Sahar Taheri, Thomas F. Rogers, Nathan Beutler, Sydney I. Ramirez, Stephen A. Rawlings, Victor Pretorius, David M. Smith, Dennis R. Burton, Laura E. Crotty Alexander, Jason Duran, Shane Crotty, Jennifer M. Dan, Soumita Das, Pradipta Ghosh

**Affiliations:** Department of Pediatrics, University of California San Diego.; Department of Computer Science and Engineering, Jacobs School of Engineering, University of California San Diego.; Moores Cancer Center, University of California San Diego.; Department of Cellular and Molecular Medicine, University of California San Diego.; Department of Immunology and Microbiology, The Scripps Research Institute, La Jolla, CA 92037, USA.; Division of Infectious Diseases, Department of Medicine, University of California, San Diego, La Jolla, CA 92037, USA.; IAVI Neutralizing Antibody Center, The Scripps Research Institute, La Jolla, CA 92037, USA.; Consortium for HIV/AIDS Vaccine Development (CHAVD), The Scripps Research Institute, La Jolla, CA 92037, USA.; Pulmonary Critical Care Section, Veterans Affairs (VA) San Diego Healthcare System, La Jolla, California; Division of Pulmonary, Critical Care and Sleep Medicine, Department of Medicine, University of California San Diego (UCSD), La Jolla, California; Center for Infectious Disease and Vaccine Research, La Jolla Institute for Immunology (LJI), La Jolla, CA, USA.; Department of Medicine, Division of Infectious Diseases and Global Public Health, University of California, San Diego (UCSD), La Jolla, CA, USA.; Department of Pathology, University of California San Diego.; Medicine, University of California San Diego.; Department of Surgery, University of California San Diego.; Division of Cardiology, Department of Internal Medicine, UC San Diego Medical Center, La Jolla 92037

**Author notes:** Co-Corresponding, Corresponding authors: Debashis Sahoo, Ph.D.; Assistant Professor, Department of Pediatrics, University of California San Diego; 9500 Gilman Drive, MC 0730, Leichtag Building 132; La Jolla, CA 92093-0831. : : Soumita Das, Ph.D.; Associate Professor, Department of Pathology, University of California San Diego; 9500 Gilman Drive, George E. Palade Bldg, Rm 256; La Jolla, CA 92093. : Pradipta Ghosh, M.D.; Professor, Departments of Medicine and Cell and Molecular Medicine, University of California San Diego; 9500 Gilman Drive (MC 0651), George E. Palade Bldg, Rm 232; La Jolla, CA 92093. Equal contribution (Debashis Sahoo, Gajanan D. Katkar).

**Keywords:** Artificial Intelligence/Machine Learning, Boolean Equivalent Clusters, Angiotensin Converting Enzyme (ACE)-2, Coronavirus COVID-19, Immune response, Lung alveoli, Natural Killer (NK) cells, Interleukin 15 (IL15)

## Abstract

We sought to define the host immune response, a.k.a, the “cytokine storm” that has been implicated in fatal COVID-19 using an AI-based approach. Over 45,000 transcriptomic datasets of viral pandemics were analyzed to extract a 166-gene signature using ACE2 as a ‘seed’ gene; ACE2 was rationalized because it encodes the receptor that facilitates the entry of SARS-CoV-2 (the virus that causes COVID-19) into host cells. Surprisingly, this 166-gene signature was conserved in all *<u>vi</u>*ral *<u>p</u>*andemics, including COVID-19, and a subset of 20-genes classified disease severity, inspiring the nomenclatures *ViP* and *severe-ViP* signatures, respectively. The *ViP* signatures pinpointed a paradoxical phenomenon wherein lung epithelial and myeloid cells mount an IL15 cytokine storm, and epithelial and NK cell senescence and apoptosis determines severity/fatality. Precise therapeutic goals were formulated and subsequently validated in high-dose SARS-CoV-2-challenged hamsters using neutralizing antibodies that abrogate SARS-CoV-2•ACE2 engagement or a directly acting antiviral agent, EIDD-2801. IL15/IL15RA were elevated in the lungs of patients with fatal disease, and plasma levels of the cytokine tracked with disease severity. Thus, the *ViP* signatures provide a quantitative and qualitative framework for titrating the immune response in viral pandemics and may serve as a powerful unbiased tool to rapidly assess disease severity and vet candidate drugs.

**One Sentence Summary:** The host immune response in COVID-19.

**PANEL: RESEARCH IN CONTEXT:** *Evidence before this study:* The SARS-CoV-2 pandemic has inspired many groups to find innovative methodologies that can help us understand the host immune response to the virus; unchecked proportions of such immune response have been implicated in fatality. We searched GEO and ArrayExpress that provided many publicly available gene expression data that objectively measure the host immune response in diverse conditions. However, challenges remain in identifying a set of host response events that are common to every condition. There are no studies that provide a reproducible assessment of prognosticators of disease severity, the host response, and therapeutic goals. Consequently, therapeutic trials for COVID-19 have seen many more ‘misses’ than ‘hits’. This work used multiple (> 45,000) gene expression datasets from GEO and ArrayExpress and analyzed them using an unbiased computational approach that relies upon fundamentals of gene expression patterns and mathematical precision when assessing them.

*Added value of this study:* This work identifies a signature that is surprisingly conserved in all viral pandemics, including Covid-19, inspiring the nomenclature ViP-signature. A subset of 20-genes classified disease severity in respiratory pandemics. The ViP signatures pinpointed the nature and source of the ‘cytokine storm’ mounted by the host. They also helped formulate precise therapeutic goals and rationalized the repurposing of FDA-approved drugs.

*Implications of all the available evidence:* The ViP signatures provide a quantitative and qualitative framework for assessing the immune response in viral pandemics when creating pre-clinical models; they serve as a powerful unbiased tool to rapidly assess disease severity and vet candidate drugs.

## INTRODUCTION

As the rapidly unfolding COVID-19 pandemic claims its victims around the world, it has also inspired the scientific community to come up with solutions that have the potential to save lives. In the works are numerous investigational drugs at various phases of clinical trials, from rationalizing^1^, to IRB approvals, recruitment and execution^2, 3^, all directed to meet an urgent and unmet need —i.e., ameliorate the severity of COVID-19 and reduce mortality.

Two obstacles make that task difficult—First, the pathophysiology of COVID-19 remains a mystery. The emerging reports generally agree that the disease has a very slow onset^4, 5^ and that those who succumb typically mount a ‘cytokine storm’^4, 6^, i.e., an overzealous immune response. Despite being implicated as a cause of mortality and morbidity in COVID-19, we know virtually nothing about what constitutes (nature, extent) or contributes to (cell or origin) such an overzealous response. Consequently, treatment goals in COVID-19 have been formulated largely as a ‘trial and error’-approach; this is reflected in the mixed results of the trials that have concluded^7^. Second, there is no established pre-clinical animal or human cell/organoid models for COVID-19; vetting the accuracy and/or the relevance of such models requires first an understanding of the host response in the disease.

We set out to define this aberrant host immune response in COVID-19 using machine learning tools that can look beyond interindividual variability to extract underlying gene expression patterns within multidimensional complex data. The approach was used across multiple cohorts of viral pandemics. The resultant pattern, i.e., signature, was subsequently exploited as a predictive model to navigate COVID-19. Findings not only pinpointed the precise nature of the cytokine storm, the culprit cell types and the organs, but also revealed disease pathophysiology, and helped formulate specific therapeutic goals for reducing disease severity. Key findings were validated in preclinical models of COVID-19 in Syrian hamsters and in the lungs and plasma of infected patients.

## METHODS

### Data Collection and Annotation

Publicly available microarray and RNA Seq databases were downloaded from the National Center for Biotechnology Information (NCBI) Gene Expression Omnibus (GEO) website^8–10^. A comprehensive catalog of these datasets is presented in **Table S1**. Gene expression summarization was performed by normalizing Affymetrix platforms by RMA (Robust Multichip Average)^11, 12^ and RNASeq platforms by computing TPM (Transcripts Per Millions)^13, 14^ values whenever normalized data were not available in GEO.

### Statistics

Boolean analysis and other statistical approaches are covered in detail in *Supplementary Online M*aterials. Briefly, the StepMiner algorithm^15^, BooleanNet statistics^16^, and BECC (Boolean Equivalent Correlated Clusters)^17^ are used to perform Boolean analyses. Gene signature is computed by using a scaled linear combination of gene expression values which is used to classify sample categories and the performance of the multi-class classification is measured by ROC AUC (Receiver Operating Characteristics Area Under The Curve) values. A color-coded bar plot is combined with a violin plot to visualize the gene signature-based classification and distribution of the gene signature score. Bubble plots of ROC-AUC values (radius of circles are based on the ROC-AUC) demonstrating the direction of gene regulation (Up, red; Down, blue) for the classification based on the 20 gene severe ViP signature and 166 gene ViP signature is visualized side by side.

### Sample size estimation

Effect size (the magnitude of the difference between groups divided by the standard deviation) for IL15 measurement is estimated as 1/0.7 from GSE157103. In order to have an 80% power (1-β = 0.8) to detect a statistically significant difference (α = 0.05) between high and low groups of patients, we need around 8 patients in each group.

### Ethics statement

Animal studies were approved and performed in accordance with Scripps Research IACUC Protocol #20-0003 (PI: Tom Rogers, PMID: 32540903). Blood from COVID-19 donors was either obtained at a UC San Diego Health clinic under the approved IRB protocols of the University of California, San Diego (UCSD; 200236X) or recruited at the La Jolla Institute under IRB approved (LJI; VD-214). COVID-19 donors were California residents, who were either referred to the study by a health care provider or self-referred. The lung specimens from the COVID 19 positive human subjects were collected using autopsy (study was IRB Exempt). All donations to this trial were obtained after telephone consent followed by written email confirmation with next of kin/power of attorney per California state law (no in-person visitation could be allowed into our COVID-19 ICU during the pandemic). The team member followed the CDC guidelines for COVID19 and the autopsy procedures^18, 19^.

### Validation studies

#### Serum cytokine measurement

Blood from COVID-19 donors was either obtained at a UC San Diego Health clinic or recruited at the La Jolla Institute under active IRB protocols. Two independent methods were used: 1) Levels of IL15 cytokine was estimated using ELISA MAX Deluxe set (Biolegend Cat. No. 435104) or Meso Scale Discovery (MSD) according to the manufacturer’s recommended protocol. 2)

#### Autopsy and biopsies of lungs

The detailed process of collecting and processing lung specimens from the COVID 19 positive human subjects is available in the supplementary methods.

#### Hamster study with anti-CoV-2 therapy

Lung samples from CoV-2-challenged 8-week old Syrian hamsters were generated exactly previously published study^20^ with either anti-spike monoclonal antibody or anti-viral therapy (EIDD-2801). See Supplementary methods for details regarding IHC protocols.

### Data deposition and materials sharing

All data is available in the main text or the supplementary materials. RNA Seq datasets from hamster studies have been deposited at NCBI GEO (GSE168095 and GSE157058).

## RESULTS AND DISCUSSION

### An ACE2-centric Study Design

To identify and validate an invariant (universal) gene signature of host response in COVID-19, we mined more than 45,000 publicly available datasets of viral pandemics across three species (human, mouse and rats) (***Step 1***; **Fig 1**). Three relatively widely accepted facts shaped our approach using Angiotensin-converting enzyme 2 (ACE2) as ‘seed’ gene in our computational studies: *(i)* ACE2 is the most well-known portal for SARS-CoV-2 entry into the host cell^21, 22^; its expression in cell lines correlates with the expression of innate immune genes ^23^ and susceptibility to SARS-CoV spike protein-driven entry^24, 25^, and its depletion in mice abrogates SARS-CoV infection^26^; *(ii)* ACE2 is a potent negative regulator of the renin–angiotensin aldosterone system (RAAS)^27^; without such restraint, the RAAS contributes to exuberant inflammation in the setting of infections^28^; and finally, *(iii)* although the mechanism through which ACE2 suppresses inflammatory response remains poorly understood, accumulating evidence indicates that infections perturb ACE2 activity, allowing for uncontrolled inflammation^29–37^.

**Figure 1.**
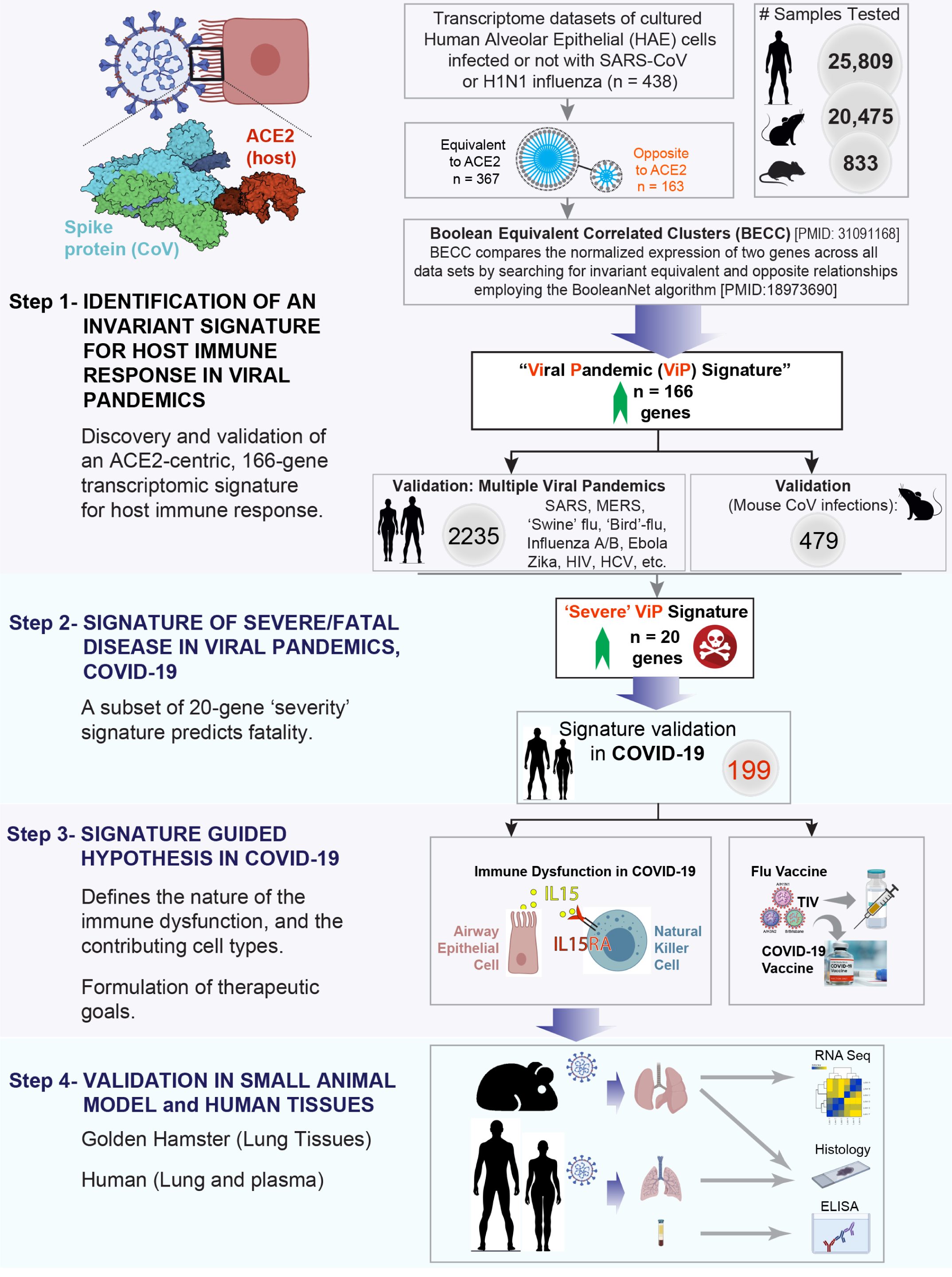
Study design. (*From top to bottom*) *<u>Step 1</u>*: A database containing > 45,000 human, mouse and rat gene– expression data was mined to identify and validate an invariant signature for host response to viral pandemic (ViP) infection. ACE2, the portal for SARS-CoV-2 entry/uptake, was used as a ‘seed’ gene and Boolean Equivalent Correlated Clusters (BECC) was used as the computational method to identify gene clusters that share invariant relationships with ACE2. Once defined, these gene clusters (a.k.a., ‘*ViP signature*’) were subsequently validated across multiple human and murine models of pandemic viral infection. *<u>Step 2</u>:* A subset of 20 genes from the ViP signature was selected that was strongly associated with severity of viral infection. These genes were validated in other cohorts to establish the ‘Severe’ ViP signature. Both 166- and 20-gene ViP signatures were validated on COVID-19 datasets. *<u>Step 3</u>*: Cross-validation studies in numerous other datasets helped-(i) define the nature (ii) and source of the cytokine storm in COVID-19, (iii) gain insights into the immunopathology of fatal disease, and (iv) set precise therapeutic goals. *<u>Step 4</u>*: Findings in step 3 were validated in hamsters and in a cohort of COVID-19 patients. A comprehensive catalog of the datasets analyzed in this work can be found in **Supplementary Table 1**.

As ***Step 2*** (**Fig 1**), we validated the signature in several human and mouse datasets of viral pandemics, and a subset of genes was identified and validated as indicators of disease severity. The signatures were then validated in SARS-CoV-2-infected cells and tissues and to explore the nature, extent and cell of origin of host response in mild and fatal COVID-19.

As ***Step 3*** (**Fig 1**), the gene signatures were prospectively used to navigate the uncharted territory of COVID-19 and pinpoint immunopathologic mechanisms, which revealed the nature (IL15), source (airway epithelium), intensity (quantitative measure) and consequence (NK cell senescence) of the cytokine storm and helped objectively formulate precise therapeutic goals to reduce the severity of COVID-19.

As ***Step 4*** (**Fig 1**), the gene signature and the mechanism of action (IL15/IL15RA) were validated in lung tissues from SARS-CoV-2 challenged golden hamster using RNASeq and IHC. In addition, precise therapeutic goal was validated in the SARS-CoV-2-challenged golden hamster model. The mechanism of action (IL15/IL15RA) was also validated by ELISA in plasma and IHC in lung tissues from UCSD COVID-19 cohort participants.

### A Shared Host Response Signature in Respiratory Viral Pandemics

Because publicly available transcriptomic datasets from SARS-CoV-2-infected samples are still relatively few, any conclusion drawn from so few samples using any computational methodology is likely to lack robustness. We chose to use an informatics approach, i.e., Boolean Equivalent Correlated Clusters (BECC)^17^, which can identify fundamental invariant (universal) gene expression relationships underlying any biological domain; in this case, we selected the biological domain of ‘*respiratory viral pandemics*’. BECC enables comparison of the normalized expression of two genes across all datasets by searching for two sparsely populated, diagonally opposite quadrants out of four possible quadrants (high-low and low-high), employing the BooleanNet algorithm^16^. There are six potential gene relationships assessed by BooleanNet: two symmetric (Equivalent and Opposite; **Fig 2A**) and four asymmetric^16^. Two genes are considered “Boolean Equivalent” if they are positively correlated with only high-high and low-low gene expression values. Two genes are considered “Boolean Opposite” if they are negatively correlated with only high-low and low-high gene expression values. Asymmetric Boolean implications result when there is only one sparsely populated quadrant. The BECC algorithm focuses exclusively on “Boolean Equivalent” relationships to identify potentially functionally related gene sets. Once identified, these invariant relationships have been shown to spur new fundamental discoveries^38, 39^, with translational potential^40^, and most importantly, offer insights that aid the navigation of uncharted territories where nothing may be known^41, 42^.

**Figure 2.**
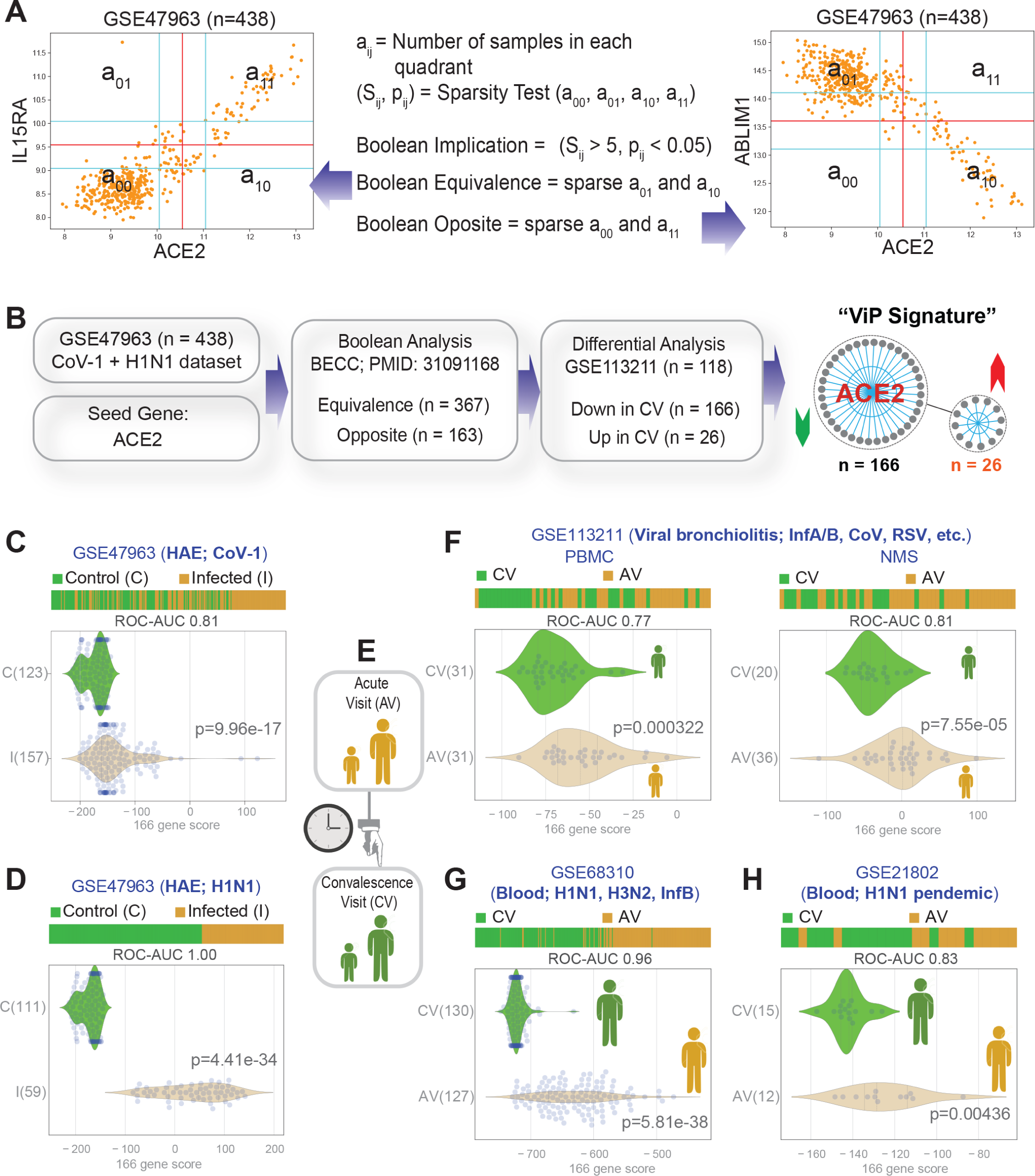
Identification and validation of an invariant ACE2-centric signature of host response to viral infections. **(A)** Computational approach to identify Boolean Equivalent and Opposite relationships. Number of samples in all four quadrants are used to compute two parameters (S, p). S > 5 and p < 0.05 is used to identify sparse quadrant. Equivalent relationships are discovered when top-left and bottom-right quadrants are sparse (*left*). Opposite relationships are discovered when top-right and bottom-left quadrants are sparse (*right*). **(B)** Schematic displaying the key computational steps and findings leading to the identification of the 166-gene host response signature using ACE2 as a ‘seed’ gene. See also **Table S2**. **(C)** Bar and violin plots displaying sample rank order (i.e., classification) of SARS-CoV-1-infected samples and distribution of the 166 gene-based signature in the test dataset (GSE47963, *in vitro* infections of human airway epithelial cells). ROC-AUC values of infected samples classifications are shown below each bar plot unless otherwise stated. **(D)** Analysis of H1N1-infected samples compared to uninfected controls using the 166-gene signature like C. **(E)** Classification of patient samples used in datasets **F-H** based on their time of collection either during ‘Acute Visit’ (AV) in the setting of an acute respiratory viral infection and ‘Convalescence Visit’ (CV) after recovery. **(F)** Analysis of PBMC samples from children (GSE113211, left) and nasal mucosal scrapings (NMS, GSE113211, right). **(G)** Analysis of peripheral blood from adults (GSE68310). **(H)** Analysis of patient samples collected during the swine flu pandemic (GSE21802).

We used GSE47963 [human airway epithelial (HAE) cultures with H1N1 and SARS-CoV infections; n = 438] as a ‘test’ dataset, which was comprised of human airway epithelial cell samples (HAE) infected *in vitro* with the causative agents of the 2009 ‘swine flu’ (influenza A-H1N1; a triple recombination of human, avian, and swine influenza viruses^43–45^) and the 2002 Severe acute respiratory syndrome (SARS-CoV-1)^46^ outbreaks (**Fig 2B**). These datasets were chosen now, and other datasets were prioritized later in the study, e.g., H5N1 (the causative agent of the avian flu in 2006-06^47^) and MERS-CoV (the causative agent of Middle East respiratory syndrome in 2012^48^) based upon the fact that they *all* contributed to outbreaks that are characterized by acute respiratory syndromes with high case-fatality rates^21^.

ACE2 is used as a ‘seed’ to identify other genes that have ‘Boolean Equivalent’ and ‘Boolean Opposite’ relationships with ACE2. These genes were subsequently filtered using differential analysis on another dataset [GSE113211 (n = 118); **Fig 2B**] that profiled heterogeneous immunophenotypes of children with viral bronchiolitis (confirmed positive for the virus in ∼100% patients; of which 25 % were infected with Influenza/Para-Influenza and 14.8% with human CoV). Transcriptomes were analyzed in nasal mucosal scrapings (NMS) and PBMC samples taken during an acute visit (AV) and during a subsequent visit at convalescence (CV)^49^. 166 genes (**Table S2**; 1-1) retained the “Boolean Equivalent” relationship with ACE2 *and* their expression was downregulated during the convalescence visit. 26 genes (**Table S2**; 2-1) retained “Boolean Opposite” relationships with ACE2 *and* their expression was upregulated during the convalescence visit. All subsequent analyses were performed using the 166 –gene signature that had Boolean Equivalent relationship with ACE2 and that was down-regulated during a convalescent visit after acute viral bronchiolitis.

First, the 166-gene signature was evaluated in the test dataset--it was used to rank order the samples and test for phenotype classification using a receiver operating characteristic curve [ROC curve; the area under this curve (AUC) represents degree or measure of separability] and displayed such classification using violin plots (**Fig 2C-D**). The signature classified the uninfected *vs*. infected samples with reasonable accuracy in the setting of SARS-CoV-1 infection (ROC-AUC = 0.81, **Fig 2C**). It also classified perfectly in the setting of H1N1 infection (ROC-AUC = 1.00, **Fig 2D**). Good classification was observed between samples from the acute visit (AV) and convalescence visit (CV) in children (test dataset; GSE113211; **Fig 2E-F**, *left*), as well as two independent adult cohorts (validation datasets that were generated in two prospective studies^50, 51^; **Fig 2G-H**). All the patients in these cohorts were infected with respiratory viruses; in one cohort, ∼45% were documented infections with pandemic Influenza strains H1N1 and H3N2 (GSE68310), whereas 100% of the patients in the other were victims of the H1N1 pandemic of 2009 (GSE21802). Regardless of the heterogeneity of these validation cohorts, the classification score using the 166-gene signature remained strong in both datasets (ROC-AUC = 0.83 - 0.96). Findings indicate that the viral pandemic signature was conserved among numerous respiratory viral pandemics, and for that reason, we christened it the ‘*Vi*ral *P*andemic’ (*ViP*) Signature.

### The *ViP* Signature Defines the ‘Cytokine Storm’ in Viral Pandemics

Reactome analyses on the 166 genes showed that the signature was largely enriched for genes within the immune system pathways, e.g., interferon and cytokine signaling, cellular processes that are critical for an innate immune response such as the ER-phagosome pathway and antigen processing and presentation, and finally the adaptive immune system (**Fig 3A-C**). In other words, the signature reflected a typical host immune response that is expected during any viral infection. This is not surprising because an overzealous host immune response, i.e., a ‘cytokine storm’ is shared among all respiratory viral pandemics (Influenza, avian and swine flu)^52^ and severe COVID-19 patients who succumb to the disease^6^. However, there were 3 surprising factors: (i) This signature and reactome profile emerged using ACE2 as a ‘seed’ gene, which is not the receptor for influenza strains to enter into host cells. (ii) It is also noteworthy that despite filtration through two unrelated datasets (**Fig 2B**), one *in vitro* and another *in vivo*, and the reduction in the number of genes in the ACE2-equivalent cluster during such iterative refinement, the pathways/processes represented in the 166-gene cluster (**Table S2**; 1-2) remained virtually unchanged. (iii) The only cytokine/receptor pair that emerged in this 166-gene cluster was interleukin-15 (IL15/IL15RA; **Fig 3A, C**), indicating that transcripts of this cytokine are invariably equivalent with ACE2 expression across all datasets analyzed. Findings are in keeping with the well-established role of IL15 in both the pathogenesis^53^ and the severity ^54^ of virus-induced lung injury. They are also consistent with the fact that IL15-/- mice are protected from lethal influenza^55^.

**Figure 3.**
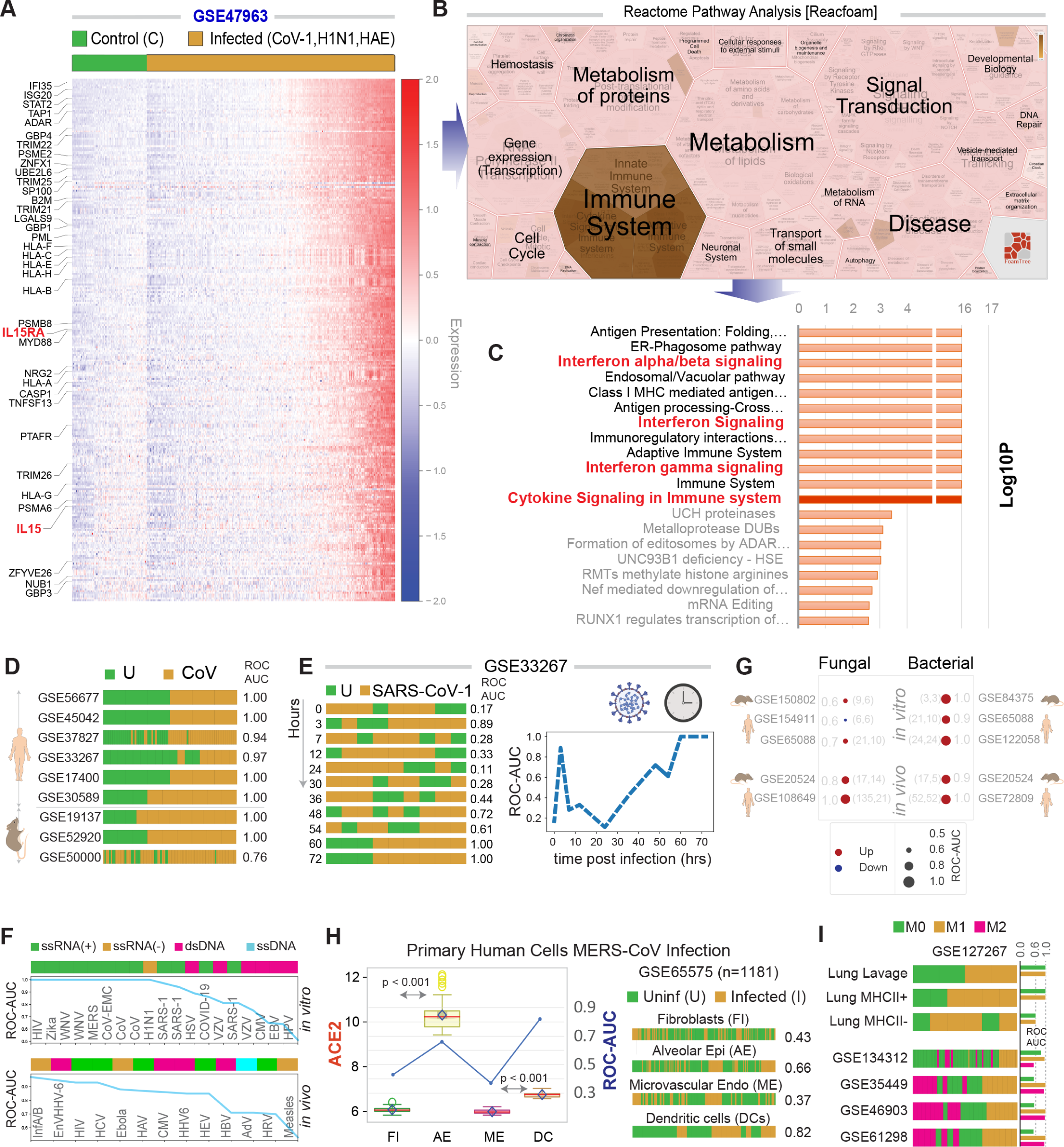
Validation of the ViP signatures in global pandemic viral infections. **(A)** Heatmap of the 166-gene signature on test dataset (GSE47963, *in vitro* infections of human airway epithelial cells). Genes that are involved in cytokine signaling in immune system are highlighted in the left. **(B)** ReacFoam analysis on the 166-gene signature that visualizes genome-wide pathway analysis based on Voronoi tessellation. **(C)** Reactome pathway analysis of the 166 genes in the Vip signature. **(D)** ViP signature-based classification of CoV-infected samples (CoV) from uninfected controls (U) in diverse human and mouse datasets. **(E)** Time course of CoV infection shows that the ViP host-response signature is slowly induced in very late (48-72 h) in Calu-3 cells infected with SARS-CoV-1. **(F)** The accuracy (Y axis; ROC AUC) of the signature to classify viral infections differs between RNA viruses and DNA viruses (*X* axis) in *in vitro* system (top). However, they are indistinguishable in *in vivo* system (bottom). See also **Table S3** and **Fig S1**. **(G)** ViP signature-based classification of human and murine samples with fungal or bacterial infections in either *in vitro* or *in vivo* settings. **(H)** The signature captures host response to CoV infection in human primary lung alveolar epithelial cells (AE) and dendritic cells (DC) better than in Fibroblasts (FI) and Endothelial (ME) cells. The accuracy of classification (ROC-AUC) strongly correlates with ACE2 expression in these cells. **(I)** Classification of macrophage polarization states ‘reactive’ (M1 polarized), unstimulated M0 and tolerant M2-like samples using the 166-gene ViP signature across diverse datasets.

Next we tested this 166-gene signature in datasets of samples infected with viruses that have either caused pandemics in the past (SARS-CoV-1, MERS, Ebola, Zika, etc.) or continue to do so at present (Influenza A/B, HIV, HCV, etc). The signature perfectly classified uninfected and infected samples (ROC-AUC = 1.00; **Fig 3D**) in four humans (GSE56677, GSE45042, GSE17400, GSE30589) and two mouse SARS-CoV1 and MERS-CoV datasets (GSE19137, GSE52920). It also performed reasonably well in two other human and one mouse datasets (ROC-AUC ranging between 0.76-0.97; GSE37827, GSE33267, GSE50000; **Fig 3D**). Analysis of a time course of infection with SARS-CoV-1 (GSE33267; **Fig 3E**) revealed that classification of infected samples improved over time, beginning at 48 h and reaching perfection (ROC AUC = 1.00) at 60-72 h, which is consistent with epidemiologic findings in prior acute respiratory viral pandemics (SARS and MERS) have average incubation periods ranging ∼2-7 days, which can sometimes last up to ∼10-14 d. Among datasets with curated samples representing other viral outbreaks and/or pandemics that are neither respiratory nor acute, we found that classification scores for RNA viruses were significantly better compared to DNA viruses in *in vitro* systems (**Fig 3F** top, **Fig S1A**), especially for those that share clathrin-dependent endocytic methods to breach host cells (**Table S3**). However, the classification scores were indistinguishable between RNA and DNA viruses in *in vivo* studies (**Fig 3F** bottom, **Fig S1A-B**). These results indicate that the 166-gene signature is shared among all viral infections, and not specific to respiratory viral pathogens.

Notably, the 166-gene host response signature was not specific for viral infections *per se*; it also performed well in classifying samples with bacterial infections, both *in vitro* and *in vivo*, and fungal infections *in vivo* (**Fig 3G**). These findings were not surprising because the prominent overrepresentation of interferon signaling that is captured within the signature (**Fig 3C**) is widely accepted as a shared fundamental aspect of host defense response during any infection^56^. Despite such apparent promiscuity, what is noteworthy is that the *ViP* signatures were relatively specific for infections/inflammation (**Fig S3**). The signature also implicated the epithelial and myeloid cells, but not ECs and fibroblasts contribute to host immune response because the classification scores were better for airway epithelial cells (AE) and dendritic cells (DC) compared to fibroblasts (FI) and microvascular endothelial cells (ME) (ROC-AUC: 0.66, 0.82 vs 0.43, 0.37; **Fig 3H**; *left*). These scores correlated well with ACE2 expression in these different cell types (p < 0.001; **Fig 3H**; *right*), raising the possibility that viral entry through the engagement of ACE2 and the induction of ACE2-equivalent host genes may be intertwined. That myeloid cells are major contributors to this signature was confirmed in five independent datasets; the 166-gene signature distinguished ‘reactive’ (M1-polarized) macrophages in them all (**Fig 3I**).

Together, these findings indicate that the ACE2-equivalent 166-gene signature is of broader relevance than just coronaviruses; the signature captures core fundamentals of host innate immune responses seen not just in respiratory viral pandemics, but viral, bacterial and even fungal infections. The airway epithelial cells and cells of myeloid lineages (DCs and macrophages) appear to be major contributors to the *ViP* signature.

### A 20-gene Subset within the *ViP* Signature Detects Disease Severity

To determine what constitutes ‘severe/fatal’ disease, we rank-ordered the 166 genes within the *ViP* signature for their ability to classify Influenza A/B-infected adult patients by clinical severity^57, 58^ (n = 154; **Fig 4A**). Severe disease was defined as intubation and mechanical ventilation due to poor oxygenation and/or death. A set of top 20 genes (**Fig 4A**; **Table S2**, **3-1; Table S4**) was sufficient to classify healthy controls from infected patients (ROC-AUC = 1.00) as well as distinguish mild from severe disease with reasonable accuracy (ROC-AUC = 0.95) in the test cohort (**Fig 4B**). Reactome pathway analyses revealed that compared to the *ViP* signature, the ‘severity’-related 20-gene cluster enriches a completely different set of cellular processes, i.e., DNA damage (especially induction of genes that are critical for base excision repair; BER), stress-induced senescence, neutrophil degranulation and changes in cell cycle (**Fig 4C, Fig S2**). We validated this signature side-by-side with the 166-gene *ViP* signature in three human datasets that included samples from mild vs. severe disease during the avian (H7N9), IAV (H3N1 and others) and the swine (H1N1) flu viral pandemics (**Fig 4D**, *left*). Both the 166-gene *ViP* signature and the 20-gene severity signature performed similarly when it came to classifying control *vs*. mild disease, but the latter performed significantly better in classifying mild *vs*. severe disease, and did so consistently in both validation datasets (ROC-AUC ranging from 0.8-0.9; **Fig 4D**, *left*).

**Figure 4.**
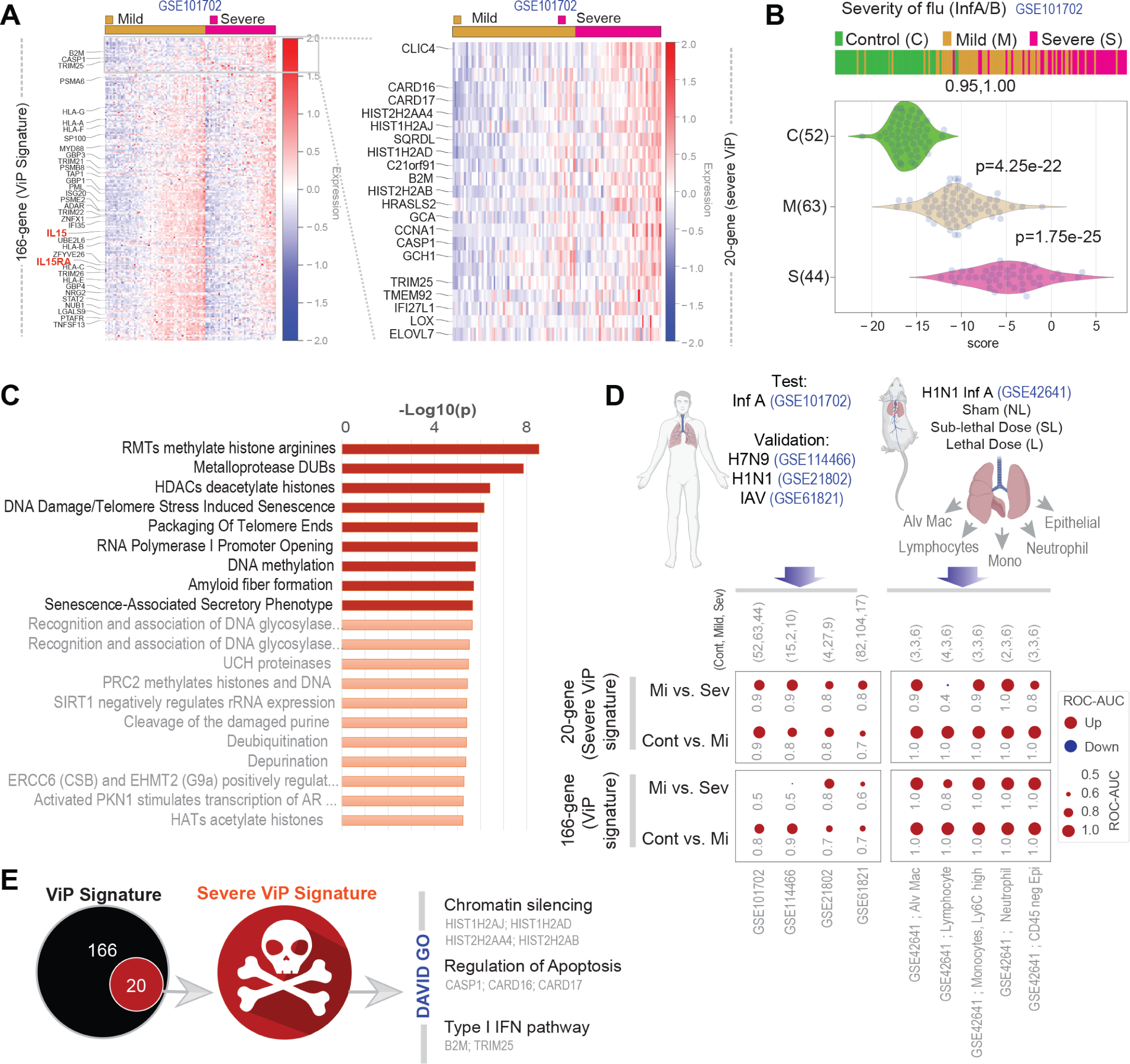
Identification of a ‘severe ViP’ signature. **(A)** Heatmap of the 166 genes on a dataset (GSE101702) annotated with varying severity of infection (healthy controls, 52; mild, 63; severe, 44). Genes are ranked based on their strength of association with severity (T-test between mild and severe). Genes that are involved in cytokine signaling in the immune system are highlighted on the left. Heatmap of top 20 selected genes (‘severe ViP’ signature) is shown on the right. **(B)** Bar and violin plots display sample rank order (i.e., classification) of patient samples and distribution of the 20-gene ‘severe ViP’ signature in the test dataset (GSE101702). ROC-AUC values of mild and severe cases are shown below the bar plot. **(C)** Reactome pathway analysis of 20 genes. **(D)** Bubble plots of ROC-AUC values (radius of circles are based on the ROC-AUC) demonstrating the direction of gene regulation (Up, red; Down, blue) for the classification based on the 20-gene severe ViP signature (top) and 166-gene ViP signature (bottom) in the test dataset (GSE101702), three more human datasets (H7N9, GSE114466; H1N1, GSE21802; IAV/H3N1 and others, GSE61821) and one mouse dataset (H1N1 Inf A, GSE42641). For each gene signature, ROC-AUC of controls vs Mild and Mild vs Severe are shown in top and bottom rows, respectively. In the mouse dataset (GSE42641) host response to lethal (L) and sublethal (SL) infection with H1N1 virus were assessed in five different lung cell types: Alv Mac, Lymphocytes, Monocytes, Neutrophil, Epithelial cells. Number of controls, mild and severe cases are shown at the top. **(E)** Summary of the 20-gene severe ViP signature and pathway analysis by DAVID GO (https://david.ncifcrf.gov/).

The severity signature performed well also in a large murine lung dataset in which mice were intranasally infected with non-lethal (NL, control), sub-lethal (SL, mild) and lethal (L, severe) doses of two different strains of H1N1 virus A; the Texas/36/91 (Tx91), which is non-lethal in C57Bl/6 mice and causes transient morbidity and compared against those infected with sublethal and lethal doses of the highly pathogenic Puerto Rico/8/34 (PR8), which causes ARDS and death in less than a week^59^. Harvested lungs were sorted into five different prospectively isolated cell subpopulations and analyzed by microarray (**Fig 4D**, *right*): alveolar macrophages, lymphocytes (BC, TC, NK), Ly6Chi mononuclear myeloid cells, neutrophils, CD45neg pulmonary epithelial cells. The 166-gene *ViP* signature distinguished the control *vs*. mild samples perfectly in all five cell types (ROC-AUC = 1.00; **Fig 4D**, *right*). The classification accuracy of the 20-gene severity signature, however, was most prominent in neutrophils (ROC-AUC = 1.00), followed by monocytes and macrophages (ROC-AUC = 0.9), and then epithelial cells (ROC-AUC = 0.8), but failed in lymphocytes. These findings suggest that the cells of the innate immune system are the primary contributors of disease severity.

We conclude that the 166-gene *ViP* signature that was initially built using *in vitro* infection datasets detects also the host immune response (‘cytokine storm’) in the complex *in vivo* systems where the response may be triggered by direct viral damage to the lung epithelium, but get propagated by feed-forward dysregulated immune response, both innate and adaptive. Surprisingly, this 166-gene *ViP* signature was not associated with disease severity; instead, severity-associated 20 genes that regulate stress and senescence-associated repression of protein expression and DNA damage (**Fig 4C**). DAVID GO analyses on the 20-gene signature indicated that 3 biological processes, e.g., transcriptional repression, apoptosis, and intermediates within the type I IFN (IFNγ signaling) pathway (**Fig 4E**) indicative of cellular distress, senescence/aging and death are the determinants of severity/fatality.

### The *ViP* Signatures are Induced in the Lung Epithelial and Immune cells in COVID-19

We next tested the ability of the *ViP* signatures to distinguish between SARS-CoV-2-infected samples and uninfected controls in 3 independent datasets, 2 of which were datasets generated from cells infected *in vitro* (**Fig 5A-C**) and one that was generated from lung samples from a fatal case of COVID-19 (**Fig 5D**). The signature perfectly classified infected from uninfected samples in them all (ROC-AUC 1.00; **Fig 5A-B-D**); of the 166 genes, both IL15 and IL15RA were notably elevated in infected samples (**Fig 5A**). The 20-gene signature performed reasonably well in distinguishing infected from uninfected A549 cells (ROC-AUC = 0.87; **Fig 5E**), and the healthy from the COVID-19 lung sample (ROC-AUC = 1.00; **Fig 5G**), but not in airway cells (bronchial; ROC-AUC = 0.57; **Fig 5F**). In fact, the 166-gene and 20-gene signatures perfectly classified infected vs. uninfected samples in all *in vitro* cellular models of CoV-2 infection, regardless of the tissue/organ (**Fig 5H***; left, middle*). The signatures performed nearly perfectly (ROC-AUC = 0.90 - 1.00; **Fig 5H**, *right*) across all lung cell types from COVID-19 infected patients analyzed by single cell sequencing.

**Figure 5.**
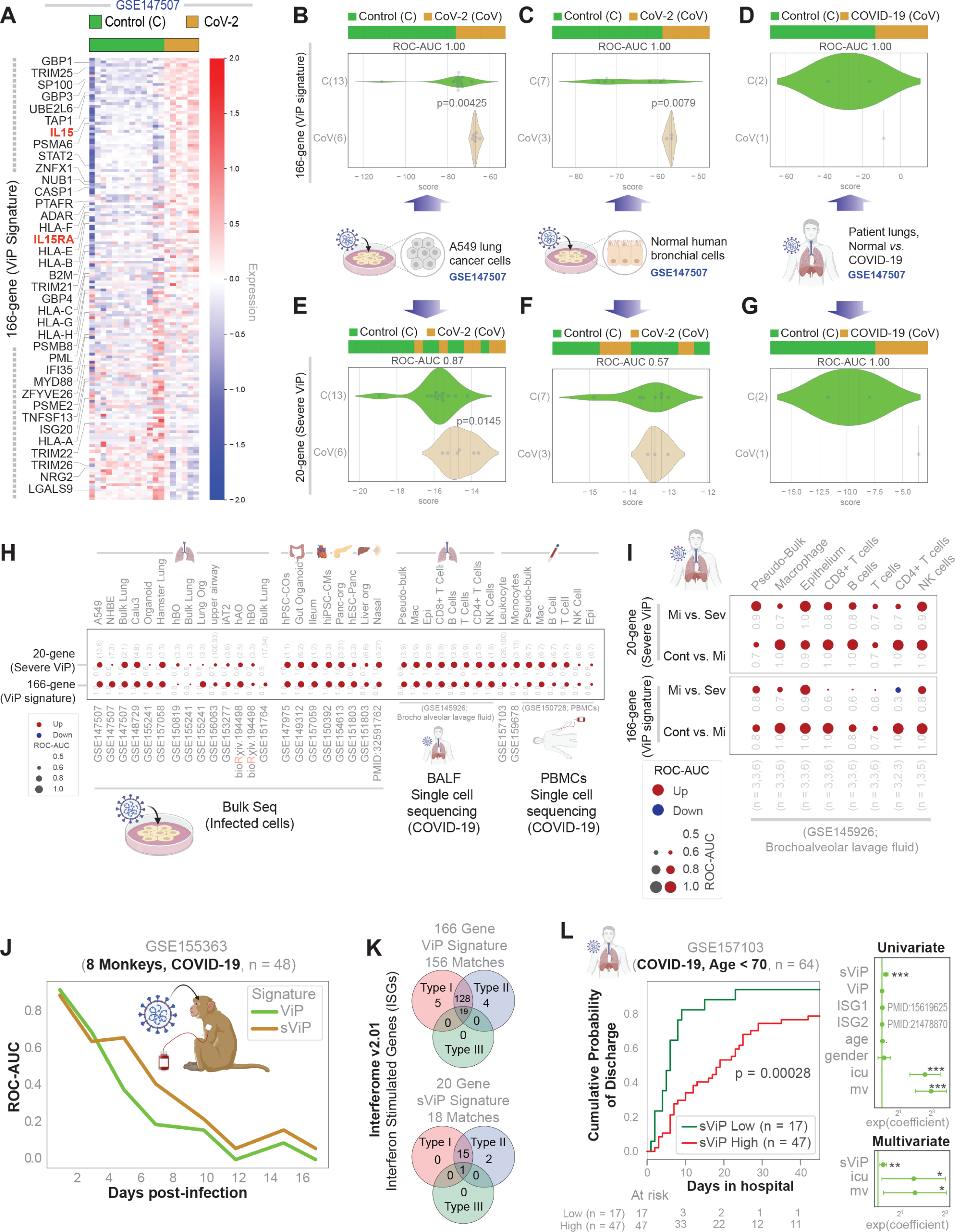
The ViP signatures define and measure the host immune response in COVID-19. **(A)** Heatmap of 166 genes in COVID-19 (GSE147507) dataset ranked by genes up-regulated in COVID-19 infected samples. Genes that are involved in cytokine signaling in the immune system are highlighted on the left. **(B-G)** Bar and violin plots displaying sample rank order (i.e., classification) and distribution of gene signature scores of COVID-19 (GSE147507) infected (CoV) and uninfected controls (C) in A549 (13 C, 6 CoV; **B, E**), normal human bronchial epithelial cells (NHBE, 7 C, 3 CoV; **C, F**), and patient lung autopsies (2 Normal, 1 CoV; **D, G**) based on 166-gene (**B, C, D**) and 20-gene ViP signatures (**E, F, G**). **(H)** Bubble plots of ROC-AUC values (radius of circles are based on the ROC-AUC) demonstrating the direction of gene regulation (Up, red; Down, blue) for the classification based on the 20 gene-severe ViP signature (top) and 166 gene ViP signature (bottom) in multiple independent datasets. **(I)** Bubble plots like panel H showing ROC-AUC of controls vs Mild and Mild vs Severe that are shown in the top and bottom rows, respectively, for each gene signature in the COVID-19 single-cell datasets (GSE145926). Dataset is analyzed as a ‘pseudo-bulk’ of all cells or after selecting individual cell types using marker genes specifically expressed in these cell types. **(J)** ViP and severe ViP (sViP) signature-based classification of blood samples (GSE155363) before and up to 17 days after COVID-19 infections in 8 monkeys using ROC-AUC measurements. **(K)** Interferon stimulated genes (ISGs) are annotated in the ViP and sViP gene lists using Interferome v2.01 web application and displayed as Venn diagrams showing the number of genes regulated by one or more IFN types (Type I, II or III). **(L)** Hospital-free days analysis (45 days followup) of COVID-19 patients (GSE157103) limited to less than 70 years old using sViP signature (low and high group) is displayed as Kaplan-Meier estimates (left) of cumulative probability of discharge and its relationship with days in hospital. Cox-proportional hazard univariate analysis (right; *top*) of sViP (high vs low) is compared to ViP signature, Interferon Stimulated Gene-signatures (ISG1^61^ and ISG2^60^), age, gender, ICU admission (icu) and mechanical ventilation (mv). Multivariate Cox-proportional hazard analysis (right; *bottom*) compares the variables that are significant in univariate settings, i.e., sViP, ICU admission (icu) and mechanical ventilation (mv).

We next tested the ability of these signatures to distinguish mild *vs*. fatal COVID-19 in single cell sequencing datasets from patient-derived lung samples (**Fig 5I**). The 166-gene signature was able to distinguish control *vs*. mild infection most effectively in macrophages, airway epithelium, CD4+ T cells and NK cells (**Fig 5I**, *lower panel, lower row*) and mild *vs*. severe disease in the epithelium and in NK cells (**Fig 5I**, *lower panel, upper row*). The 20-gene signature not only performed well in classifying control *vs*. mild infection in the same 4 cell types as above but also in B cells and CD8+ T cells (**Fig 5I**, *upper panel, lower row*). However, the 20-gene severity signature continued to perform most optimally in the epithelium (ROC-AUC = 1.00) and in NK cells (**Fig 5I**, *upper panel, upper row*). The signatures were rapidly induced also in monkeys challenged with SARS-CoV-2, and gradually suppressed during convalescence after 17 d (**Fig 5J**). Because the ViP signature is comprised of IFN-signaling pathways and presumably IFN-stimulated genes (ISGs), we asked if the ViP/severe-ViP signatures offer any additional advantage beyond ISGs. Using Interferome v2.01 (http://www.interferome.org) we first confirmed that 155/166 genes in ViP signature and 18/20 genes in severe-ViP signatures were genes that are likely to be regulated by IFN signaling (**Fig 5K**). Surprisingly, despite such high degree of pathway overlap with ISGs, the severe-ViP signature (sViP) was able to prognosticate outcome (hospital-free days) in a cohort of patients with COVID-19 (**Fig 5L**; *left*). When compared head to head in an univariate analysis using Cox proportional hazards regression model, the prognostic effect of the severe-ViP signature emerged as superior to two different sets of previously published ISGs^60, 61^ in their ability to prognosticate hospital-free days (**Fig 5L**; *top right*). Three factors emerged as determinants of longer hospital stays: (i) the ICU admission status, (ii) need for mechanical ventilation and (iii) induction of the severe-ViP signature. A multivariate analysis using Cox proportional hazards regression model suggested that these three factors may be independent covariates of poor outcome (**Fig 5L**; *bottom right*).

Together, these findings show that the 166-gene *ViP* signature seen in other respiratory viral pandemics is conserved also in COVID-19. The cytokine storm (166-genes, which included IL15/IL15RA; **Table S2**) was induced in multiple cell types; however, the 20-gene *ViP* signature of disease severity and fatality was most prominently induced in two cell types: (i) the airway epithelial cells, known producers of IL15 after viral infections^62, 63^ and (ii) the NK cells which are known targets of physiologic as well as overzealous IL15 response^64, 65^. Findings also show that despite the ISG-like makeup of the ViP signatures, there are key components within the signature that is able to detect disease severity.

### Viral infection and IL15 Induce, and Flu Vaccine Attenuates the *ViP* Signatures in NK cells

We next asked how the *ViP* signatures are impacted when NK cells are exposed to virus-infected epithelial cells. NK cells are known to lyse influenza virus-infected cells by direct cytotoxicity and antibody-dependent cellular cytotoxicity (ADCC); enhancing such NK cell function has been shown to control influenza virus infections ^66^. Clearance of other viruses (HIV-1, other retroviruses, etc.) and cancer immunotherapies also leverage such NK cell-dependent ADCC^67, 68^. We analyzed a transcriptomic dataset (GSE115203)^69^ generated from co-culture studies of human PBMCs (3 donors) with influenza (H1N1 Puerto Rico/08/1934)-infected airway epithelial cells (A549) (**Fig 6A**; *top*). PBMCs (from co-culture), or NK cells FACS-sorted from the PBMC were then analyzed by RNA Seq, and the study had confirmed NK cell ADCC responses were durably induced in this assay *via* type I IFN release from PBMCs. We found that both the 166- and 20-gene *ViP* signatures were induced in PBMCs and in NK cells sorted from the PBMCs (**Fig 6A**; *bottom left*), indicating that NK cells in these co-culture models were sufficient to capture the observed host immune response in patients with COVID-19.

**Figure 6.**
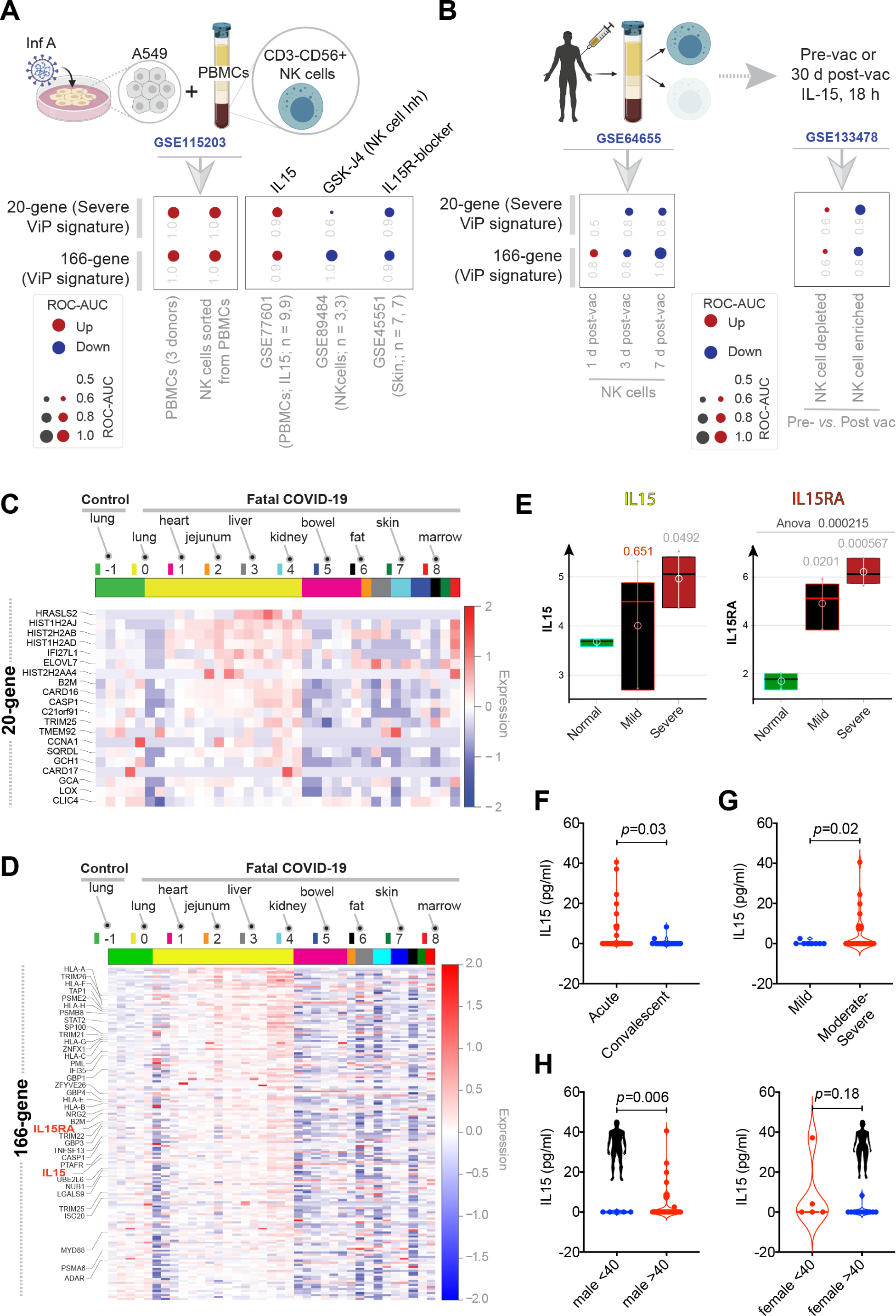
ViP signatures reveal an interplay between IL15-storm and NK cell dysfunction in fatal COVID-19. **(A)** Bubble plots of ROC-AUC values (radius of circles are based on the ROC-AUC) demonstrating the direction of gene regulation (Up, red; Down, blue) for the classification based on the 20 gene severe ViP signature (top) and 166 gene ViP signature (bottom) in following datasets. RNASeq data (GSE115203) from PBMCs and sorted NK cells from PBMCs incubated with uninfected A549 cells for 12 hrs compared to infected A549 cells. PBMCs treated with IL15 compared to IL2 (GSE77601). RNASeq analysis of NK cells (GSE89484) treated with GSK-J4 compared to DMSO. Skin tissue in mice (GSE45551) is treated with anti-IL15RB antibody compared to PBS. **(B)** RNASeq data of NK cells isolated from two donors prior to vaccination compared (left) to days 1, 3, and 7 post-TIV vaccination like panel A. RNASeq data of NK enriched and NK depleted PBMCs from healthy donors compared to 30 day post-vaccination like panel A. **(C, D)** Heatmap of 20-gene (panel **C**) and 166-gene (panel **D**) ViP signatures in tissues collected during rapid autopsies on patients who succumbed to COVID-19. Genes are ranked according to the strength of differential expression (T-test) in lung tissue between normal and infected tissue. **(E)** Box plots of IL15 and IL15RA in samples from varying severity of COVID-19. **(F-H)** Violin plots show levels of plasma IL15 in COVID-19 patients stratified by disease acquity (F), by clinical severity (G) and by gender and age (H). Welch’s two sample unpaired t-test is performed to compute the p values. See also **Table S5** for patient metadata.

To test the role of IL15 in the induction of *ViP* signatures, we leveraged three datasets— one that used recombinant IL15 (PBMCs; GSE77601), another that used anti-IL1Rβ mAb (mouse skin biopsies; GSE45551)^70^, and a third study using the prototypic H3K27 demethylase inhibitor, GSK-J4; the latter was shown to inhibit NK cell effector cytokines in response to IL15 without impacting its cytotoxic killing activities (human, NK cells; GSE89484)^71^. Both *ViP* signatures were stimulated by IL15, but attenuated in the two other datasets where IL15’s actions were blocked pharmacologically (**Fig 6A**; *right*). These findings indicate that IL15 could be necessary and sufficient to induce the *ViP* signatures.

Because two independent studies^72, 73^ recently showed that those vaccinated against influenza have lower odds of requiring intensive care, invasive ventilation and/or dying, we analyzed two transcriptomic datasets (GSE64655^74^ and GSE133478^75^) in which PBMCs from subjects vaccinated with seasonal trivalent or quadrivalent influenza vaccine (TIV/QIV) were collected and analyzed for NK cell activation. The first study showed that the NK cells continued to demonstrate progressive attenuation of both the 166- and 20-gene signatures rapidly within 7 days (**Fig 6B**, *left*). The second study, in which the NK cell-enriched and depleted fractions collected pre- and post(30 d)-vaccination were tested for their response to re-stimulation with IL15 (low dose, 0.75 ng/ml, 18 h); such stimulation is known to enhance NK cell activity ^76–78^ and promote viral clearance ^79–81^. Both ViP signatures were attenuated post-vaccination in NK cell-enriched fractions, but not in depleted fractions (**Fig 6B**, *right*). Because such post-vaccination attenuation happened in the setting of experimentally confirmed^75^ enhancement of overall NK cell response, we conclude that attenuation of *ViP* signatures among recipients of TIV could continue to offer protection during re-challenge. Because such protection is seen in NK-cell enriched, but not depleted fractions, we conclude that the protection is mediated primarily *via* the preservation of functional NK cells.

### An IL15-storm Originating in the Lung Alveoli Determines the Severity of COVID-19

We next analyzed the *ViP* signatures in transcriptomic datasets generated from multiple organs at autopsy. Both the 166- and 20-gene *ViP* signatures were predominantly enhanced in one organ, the lungs (**Fig 6C-D**); and IL15/IL15RA were also elevated in the lungs (**Fig 6D**). These findings indicate that the 20- and 166-gene signatures go together and suggest a plausible cause and effect relationship. For instance, severity-related cellular events (such as epithelial and NK cell senescence) occur in the milieu of the organ that mounts the highest IL15-predominant cytokine response, i.e., lungs. We also found that IL15 and its receptor IL15RA were significantly increased in severe COVID-19 lungs (**Fig 6E**). These findings predict that an overzealous IL15-predominant cytokine response is the most consistent finding in the most severe cases of COVID-19 and that the lung epithelium is the likely source of such a storm.

These predictions were validated in a cohort of symptomatic COVID-19 patients who presented to the UC San Diego Medical Center with varying disease severity, ranging from mild to fatal (see **Table S5**). Plasma ELISA studies revealed that IL15 levels were significantly elevated during the acute compared to the convalescent visit (**Fig 6F**), and in whom the clinical presentation was moderate-to-severe compared to those with mild disease (**Fig 6G**). A sub-group analysis confirmed that while gender or age did not have a significant impact on plasma IL15 levels independently, the aged male (> 40 y) cohort had a significantly higher IL15 level than the young males (**Fig 6H**; *left*). No such pattern was noted among females. These findings are consistent with the fact that the gender gap in COVID-19-related deaths widens markedly with age^82^. Lungs collected during autopsies from patients who succumbed to COVID-19 (see **Table S6**) further confirmed that lung epithelial cells, especially the alveolar type II pneumocytes and alveolar immune cell infiltrates express high levels of IL15 and its receptor, IL15RA (**Fig 7A-B**).

**Figure 7.**
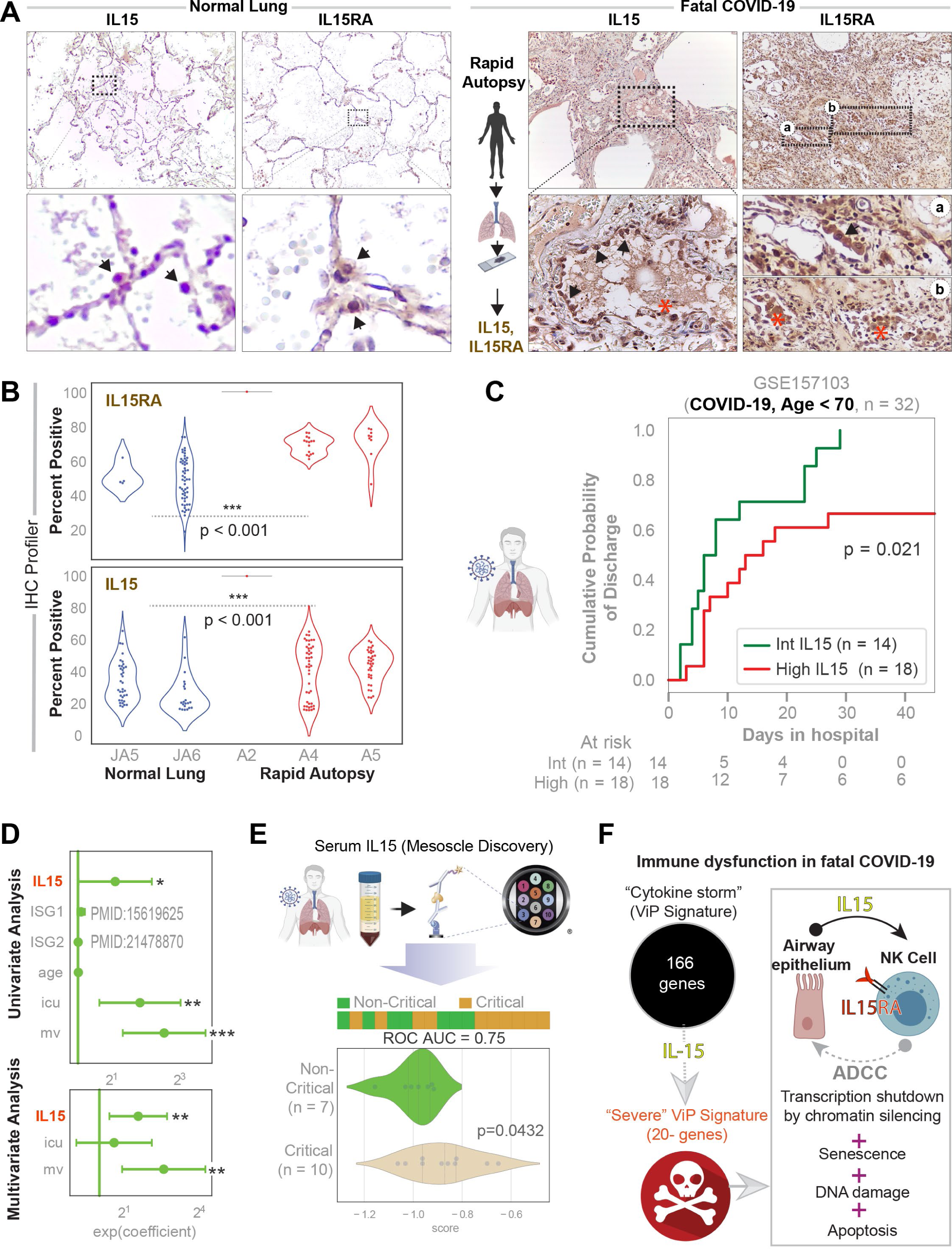
Lung alveolar cells contribute to the IL15 storm in fatal COVID-19. (**A**) Normal lung tissue obtained during surgical resection (*left*) or lung tissue obtained during autopsy studies on COVID-19 patients (*right*) were stained for IL15 and IL15RA. Representative images are shown. Mag = 10X. (**B**) Violin plots display the intensity of staining for IL15RA (top) and IL15 (bottom), as determined by IHC profiler. (**C**) Hospital-free days analysis (45 days followup) of COVID-19 patients (GSE157103) limited to males less than 70 years old using the abundance of IL15 transcripts (intermediate and high groups) is displayed as Kaplan-Meier estimates (left) of cumulative probability of discharge and its relationship with days in hospital. (**D**) Cox-proportional hazard univariate analysis (right; *top*) of sViP (high vs low) is compared to ViP signature, Interferon Stimulated Gene-signatures (ISG1, PMID:15619625; ISG2, PMID:21478870), age, gender, ICU admission (icu) and mechanical ventilation (mv). Multivariate Cox-proportional hazard analysis (right; *bottom*) compares the variables that are significant in univariate settings, i.e., sViP, ICU admission (icu) and mechanical ventilation (mv). (**E**) *Top*: Schematic displays the workflow for patient blood collection and assessment of IL15 levels by mesoscale. *Bottom*: Bar (top) and violin (bottom) plots for the levels of IL15 cytokine (score = Z score of the log reduced mesoscale concentration data). ROC AUC numbers indicate the strength of classification between patients with critical/fatal disease course vs. those with non-critical infection. (**F**) Summary of IL15 signaling and the hypothetical role of NK cells in the severity of COVID-19 infections.

Finally, in a cohort of patients with COVID-19, high levels of IL15 transcript carried a poor prognosis (lower probability of discharge from the hospital; **Fig 7C**). An univariate analysis using Cox proportional hazards regression model showed that the prognostic effect of high-IL15 was superior to ISGs^60, 61^ (**Fig 7D**; *left*), as we observed previously for ViP signatures (**Fig 5L**). A multivariate analysis using Cox proportional hazards regression model suggested that need for mechanical ventilation and IL15 induction may be independent covariates of poor outcome (**Fig 7D**; *right*). That the serum IL15 levels track disease severity was validated in a cohort of patients presenting to our institution with a diagnosis of COVID-19; critical/fatal disease was associated with significant elevation of the cytokine (**Fig 7E**).

Taken together, these findings support the following model of the immunopathogenesis of COVID-19 (**Fig 7C**): Airway epithelial cells and cells of the myeloid lineage and other immune cells are the primary source of the 166-gene cytokine storm, of which, IL15 is a component. It is possible, that the primary target of IL15, i.e., NK cells, when exposed to this storm for a prolonged period undergo damage, stress-induced senescence and apoptosis. Our model is consistent with prior studies showing that the airway epithelial cells (especially bronchial) constitutively express the IL15 and IL15RA/B genes and that viral infections^63^ and IFNγ induced the synthesis and secretion of IL15^63^ and that prolonged and excessive stimulation with IL15 is known to induce NK cell exhaustion ^64, 65^. These findings are consistent with the emerging reports that NK cells are significantly exhausted and reduced in cases of severe COVID-19 infection^83, 84^ and that such reduction was seen as early as 3-6 days after the onset of symptoms^85^. We conclude that fatal COVID-19 is characterized by a paradoxical immune response, i.e., suppression of epithelial and NK cell functions (immunosuppression) in the setting of a cytokine storm (overzealous immune response).

### The ViP Signatures Formulate Therapeutic Goals, Track Treatment Efficacy

Previously we showed that the attenuation of the *ViP* signature was ‘associated’ with the acquisition of natural convalescence in several respiratory viral pandemics (**Fig 2F-H**); we now asked if they could serve as a readout of therapeutic efficacy. We analyzed interventional studies in the setting of other viral infections that shared the *ViP* signature, i.e., HCV, HIV, Zika and Ebola (**Fig 3H**; **Fig S1**; **Table S3**). The 166-gene *ViP* signature classified HCV-infected liver biopsies treated or not with directly acting anti-viral agents (DAAs) (**Fig 8A-C**) and HIV-infected samples treated or not with *<u>a</u>*nti-*<u>r</u>*etroviral *<u>t</u>*herapeutics (ART; ROC-AUC = 1.00; **Fig 8D**) with sufficient accuracy. In the case of Ebola, the *ViP* signature was somewhat effective in classifying crisis (i.e., acute) from convalescent PBMC samples (ROC AUC 0.64; **Fig S4A, top**), and previously described anti-Ebola therapeutic strategies (Topoisomerase depletion with siRNA^86^ inhibited the signature in Ebola-infected alveolar epithelial cells (siTop; ROC AUC 1.00; **Fig S4A, bottom**)^86^. Finally, the *ViP* signature was accentuated in Zika infected human cortical neural progenitor cells (**Fig S4B**) and was effectively attenuated when these infected samples were treated with two investigational drugs that was found to be effective in inhibiting Zika infection. These findings imply that attenuation of the 166-gene *ViP* signature is a desirable therapeutic goal.

**Figure 8.**
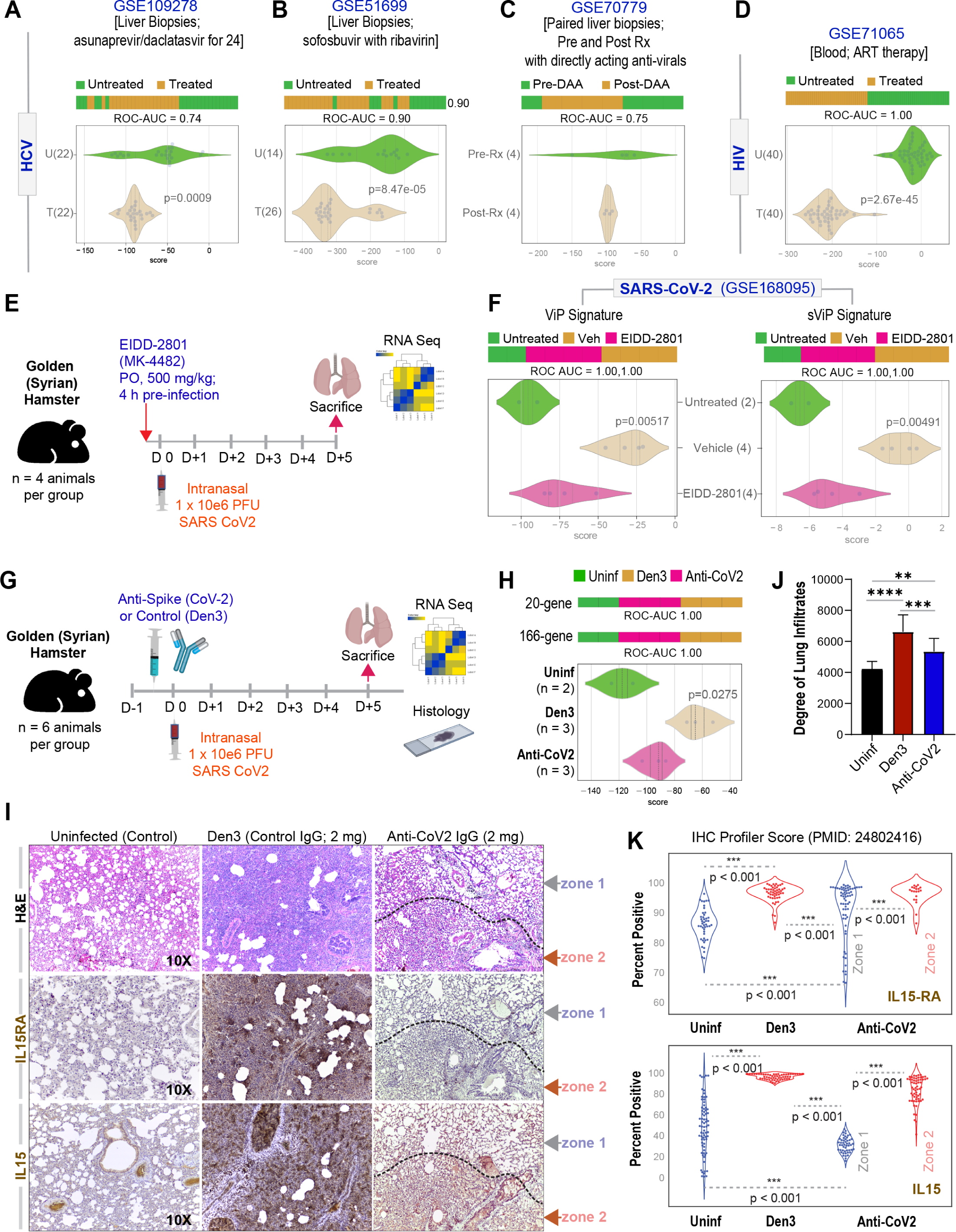
Validation of ViP signature-guided therapeutic goals. **(A-C)** The 166-gene ViP signature-was used to classify liver biopsies from HCV-infected patients treated or not with directly acting anti-viral agents. ROC-AUC values are shown below each bar plot unless otherwise stated. **(D)** 166-gene ViP signature-based classification of blood samples from HIV-infected patients treated with anti-retroviral therapy (ART). **(E)** The compound EIDD-2801 (MK-4482; 500 mg/kg) or vehicle (*Veh*) was administered at indicated doses to Golden Syrian hamsters 4 h prior to intranasal infection with SARS-CoV-2. Hamsters were were sacrificed on day 5 and lungs we analyzed by RNA sequencing. **(F)** Bar (top) and violin (bottom) plots using the ViP (left) or sViP (right) signature-based classification of lung samples from hamsters in E and uninfected controls. **(G)** Schematic showing the experimental design for validating the ViP signatures as useful tools to assess therapeutic efficacy. *Uninf*, uninfected; *Den3 and Anti-CoV-2* indicate SARS-CoV-2 challenged groups that received either a control mAb or the clone CC12.2 of anti-CoV-2 IgG, respectively. (**H**) Bar (*top*) and violin (*bottom*) plots display the 166- and 20-gene ViP signatures in the uninfected and the SARS-CoV-2 challenged groups, treated with control or anti-CoV-2 IgG. (**I-K**) Lungs harvested from the 3 groups of hamsters were analyzed by H&E and IHC. Representative images are shown in I. Mag = 10X. Bar graphs in J display the abundance of cellularity and infiltrates in the lungs of the 3 groups, as determined by ImageJ. Violin plots in K display the intensity of staining for IL15RA (*top*) and IL15 (*bottom*), as determined by IHC profiler.

We next sought to determine if the SARS-CoV-2 virus can induce the *ViP* signatures, and whether the signatures can track therapeutic response. We tested two therapeutic approaches. The first approach was the use of N-hydroxycytidine, the parent of the prodrug MK-4482 (Molnupiravir, EIDD-2801) which has not only proven as a potent and selective oral antiviral nucleoside analogue in mice, guinea pigs, ferrets and human airway epithelium organoids^87–92^, but also showing promise in Phase IIa trials in the treatment of COVID-19 patients (NCT04405570). We analyzed by RNA seq the lungs from SARS-CoV-2-challenged golden Syrian hamsters who were pre-treated either with this drug or vehicle control (see study protocol in **Fig 8E**). Both 166- and 20-gene ViP signatures were induced in the vehicle-treated arm, and effectively suppressed in the drug-treated arm to levels close to uninfected controls (GSE168095; **Fig 8F**). The second approach was the use of SARS-CoV-2-neutralizing antibodies whose design was inspired by monoclonal antibodies (mAbs) isolated from from convalescent donors^20^. A specific isotype of this antibody, which binds to the receptor-binding domain (RBD-A) of SARS-CoV-2 spike protein in a fashion that precludes binding to host ACE2, was demonstrated as effective in preventing infection and weight loss symptoms, in cell-based and *in vivo* hamster models of infection, respectively. We observed that SARS-CoV-2-challenged hamsters that were pre-treated with anti-CoV-2 antibody, but not the control Den3 antibody (see **Fig 8H** for study protocol) had 3 key findings: (i) they suppressed both the 166- and 20-*ViP* signatures that were otherwise induced in the infected lungs (GSE157058; **Fig 8I**); (ii) their lungs were protected from overwhelming immune cell infiltration and obliteration of alveolar space (**Fig 8J-K**); (iii) expression of IL15 and IL15 receptor was significantly reduced compared to what was observed in the infected lungs (**Fig 8J, L**).

These results validate the ACE-centric computational approach for identifying the *ViP* signatures, i.e., when ACE2•virus engagement was disrupted using antibodies, or reduced using directly acting anti-virals that prevent viral replication using Molnupiravir, the signatures were suppressed. The findings also indicate that the reversal of the signature and the IL15 storm could be used as a readout of therapeutic efficacy.

## CONCLUSION

The major and unexpected finding in this work is that all viral pandemics (regardless of their acuity, causative virus, case fatality rates and clinical presentation) share a common fundamental host immune response. Summarized below are our three major findings.

*First,* we defined an invariant 166-gene host response --the so-called “cytokine storm”– that is surprisingly conserved among all viral pandemics. What was also unusual is that the signature emerged despite the rationalized use of ACE2 as a ‘seed’ gene to identify ‘coronaviral infection-associated genes’. This suggests that while ACE2 may be the entry site for SARS-CoV-2, it is a prominently upregulated gene during host response to other viral infections. As a key regulator of the renin-angiotensin system (RAS), ACE2 expression is increased in the setting of multiple stressors, including non-CoV-2 infections. For example, IAV, H7N9 and rhinoviruses amplify the expression of ACE2 in the distal lung^93–95^. ACE2 activity is induced also in bacterial lung infections^29^. In fact, ACE2 protects against acute lung injury in several animal models of ARDS^96^. Thus, retrospectively, ACE2 is not as specific a ‘seed’ gene for SARS-CoV-2 as was assumed, and hence, it is not so surprising that the ACE2-equivalent ViP signature is more generalizable as a signature that is induced in respiratory infections. By comparing the *ViP* signatures head-to-head against ISGs, we presented evidence that *ViP* signatures, but not ISGs have prognostic significance. Regardless, our findings show that the ViP signatures capture the quality/nature of the shared ‘invariant’ host response, demonstrating their potential usefulness as diagnostic tools. The signatures also provide a ‘quantitative’ framework for assessing the variable degrees of such host response across patients/conditions, reflecting variable degrees of morbidity and mortality across the pandemics/outbreaks; these findings demonstrate their potential usefulness as prognostic tools. We conclude that the ViP signatures not just define the nature of the host immune response to viral pandemics, but also allows the tracking and quantification of such response.

*Second,* we define the precise nature of the cytokine storm and pinpoint the IL15 cytokine and its receptor, IL15RA as an invariant component. We demonstrate that systemic levels of IL15 tracks with disease severity among patients and that the levels are notably elevated in the aged male (the predisposed age group in COVID-19, as per reports worldwide). Using a combination of single cell RNA Seq and human lung histology, we also pinpoint the lung epithelial and myeloid cells as the key contributors to the *ViP* signature, and more specifically, IL15/IL15RA. These findings were recently validated in another concurrent publication^97^--multivariate analyses of soluble biomarkers identified that increased IL-15 is independently associated with mortality and that the levels of the cytokine was consistently high throughout the hospitalization in patients who died versus those who recovered.

*Third*, we found that a subset of 20-gene ‘severe’ *ViP* signature, indicative of stress-induced senescence, transcriptional repression, DNA damage and apoptosis is also shared among various viral pandemics. In patients with COVID-19, this signature was seen in lung epithelial and NK cells, which is intriguing because airway epithelial cells is a prominent source and the NK cells are a major target of IL15. Thus, the *ViP* signatures begin to paint a picture of ‘paradoxical immunosuppression’ at the heart of fatal COVID-19, in which, the observed NK cell exhaustion/depletion in severe COVID-19^83–85, 98^ could be a consequence of an overzealous IL15 storm, leading to their senescense and apoptosis.

In closing, given that the emerging pandemic is still largely a mystery to us in terms of how it picks its victims, the *ViP* signature we define here provides a computational framework for navigation in otherwise uncharted territory. While it is expected that the signature will be more effective and accurate when it is iteratively filtered using emerging COVID-19 datasets, we provide evidence for its usefulness now in formulating therapeutic strategies and rapidly screening for therapeutics. Because the *ViP* signature of host response is seen also in other viral pandemics tested, findings may also be relevant also in navigating management strategies in those pandemics.

## ACKNOWLEDGEMENTS

This work was supported by the National Institutes for Health (NIH) grants R00-CA151673 and R01-GM138385 (to DS) and R01-AI141630, CA100768 and CA160911 (to P.G), R01DK107585 (SD) and R01-AI155696 (to P.G, D.S and S.D), UCOP-RGPO (R00RG2628 & R00RG2642 to P.G, D.S and S.D), The Sanford Stem Cell Clinical Center at UCSD (to P.G, D.S and S.D) and U19-AI142742 (to S.C, CCHI: Cooperative Centers for Human Immunology), and LJI Institutional Funds (to S.C). Dr. Crotty Alexander’s salary was supported in part by the VA San Diego Healthcare System. The authors would like to thank Rachel White (RW) and Jen Bigbee (JB), who assisted with both thoracotomies/biopsies and helped set up and design the autopsy study.

## AUTHOR CONTRIBUTIONS

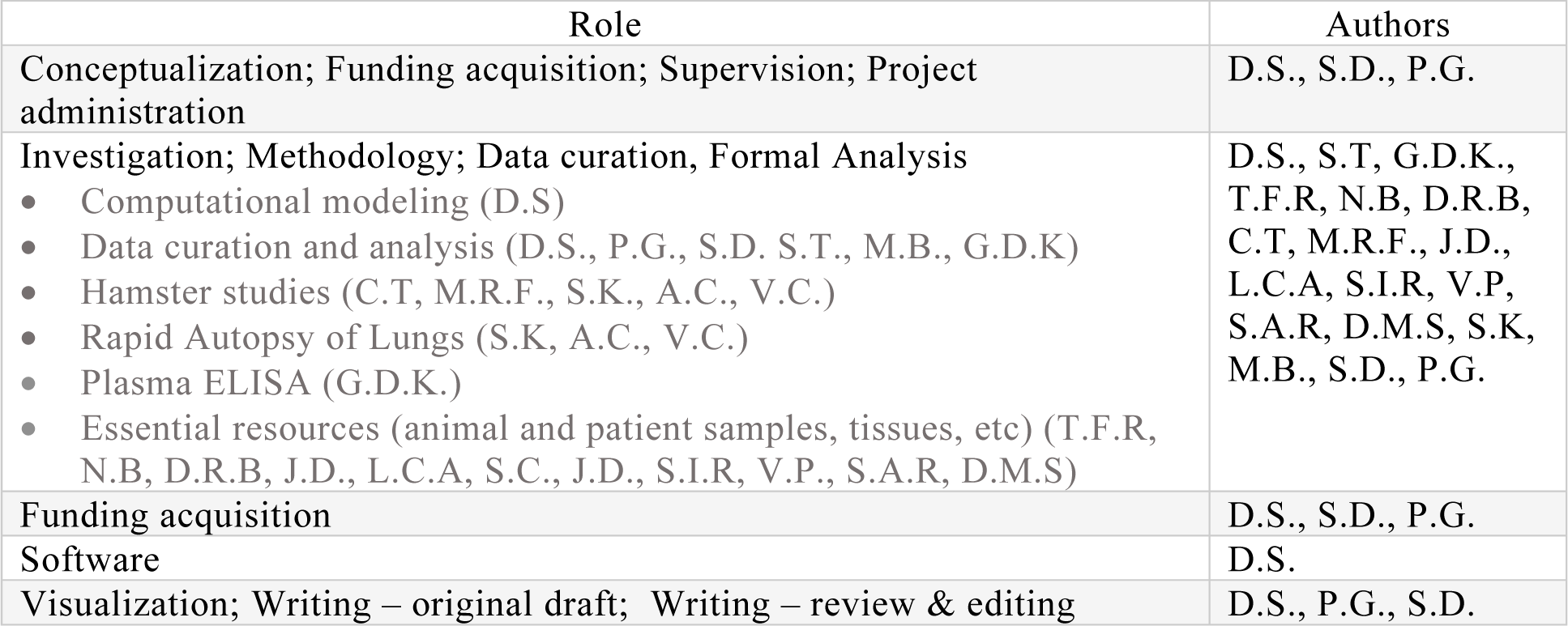

## Supplementary Materials: Includes

- Detailed Materials and Methods
- Supplementary Text-**n/a**
- Figures **S1, S2**
- Tables **S1-6**
- External Databases **-None**

---

**Table S1.**
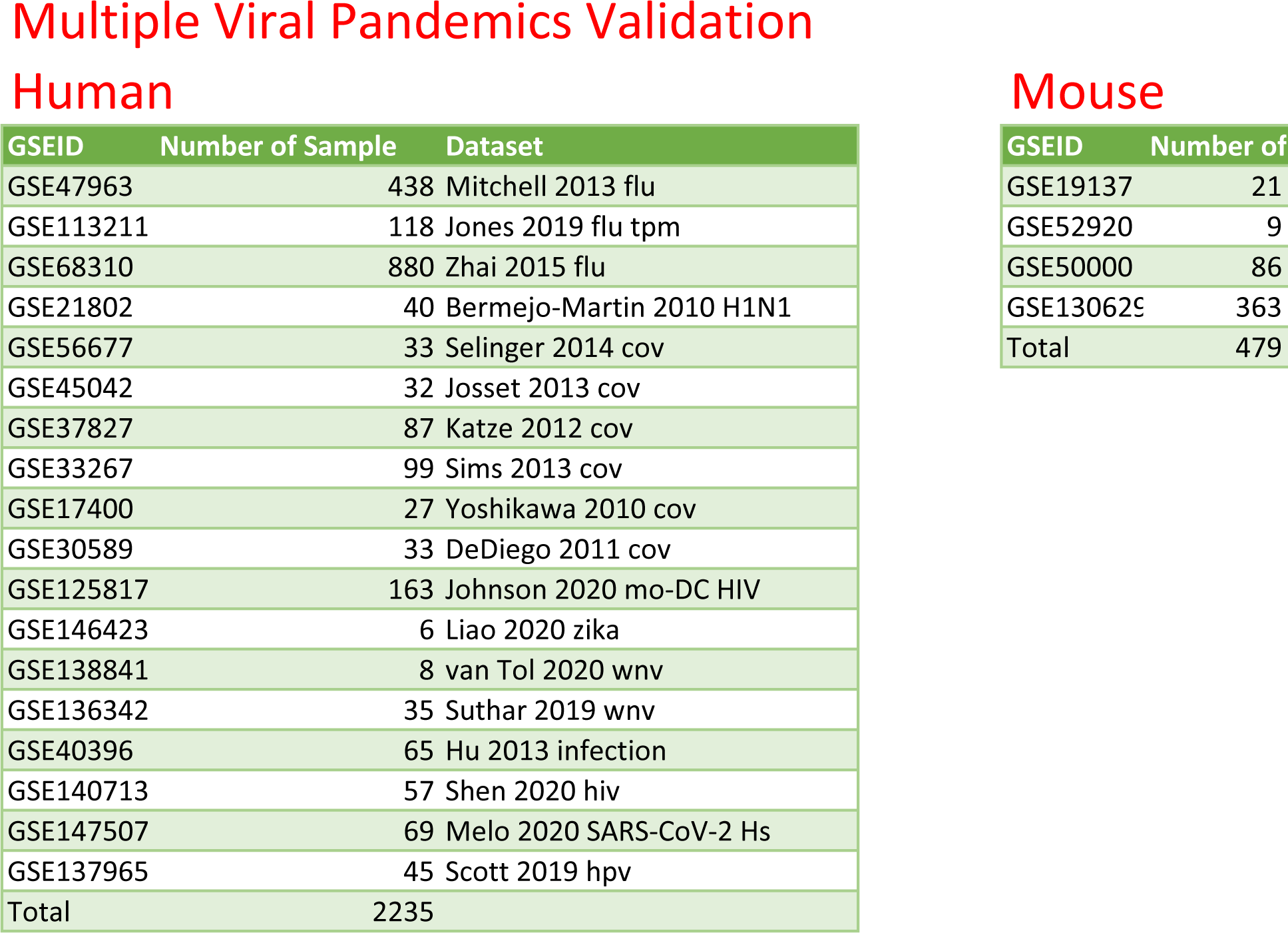
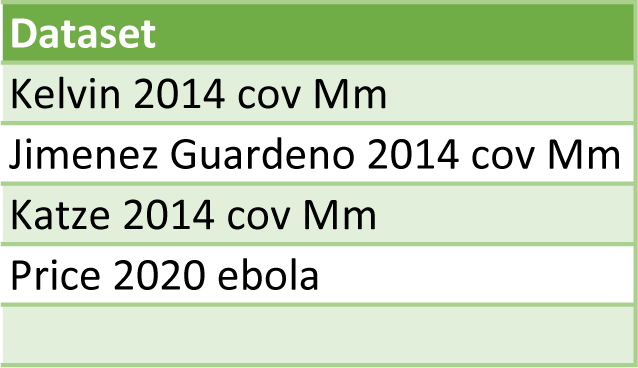
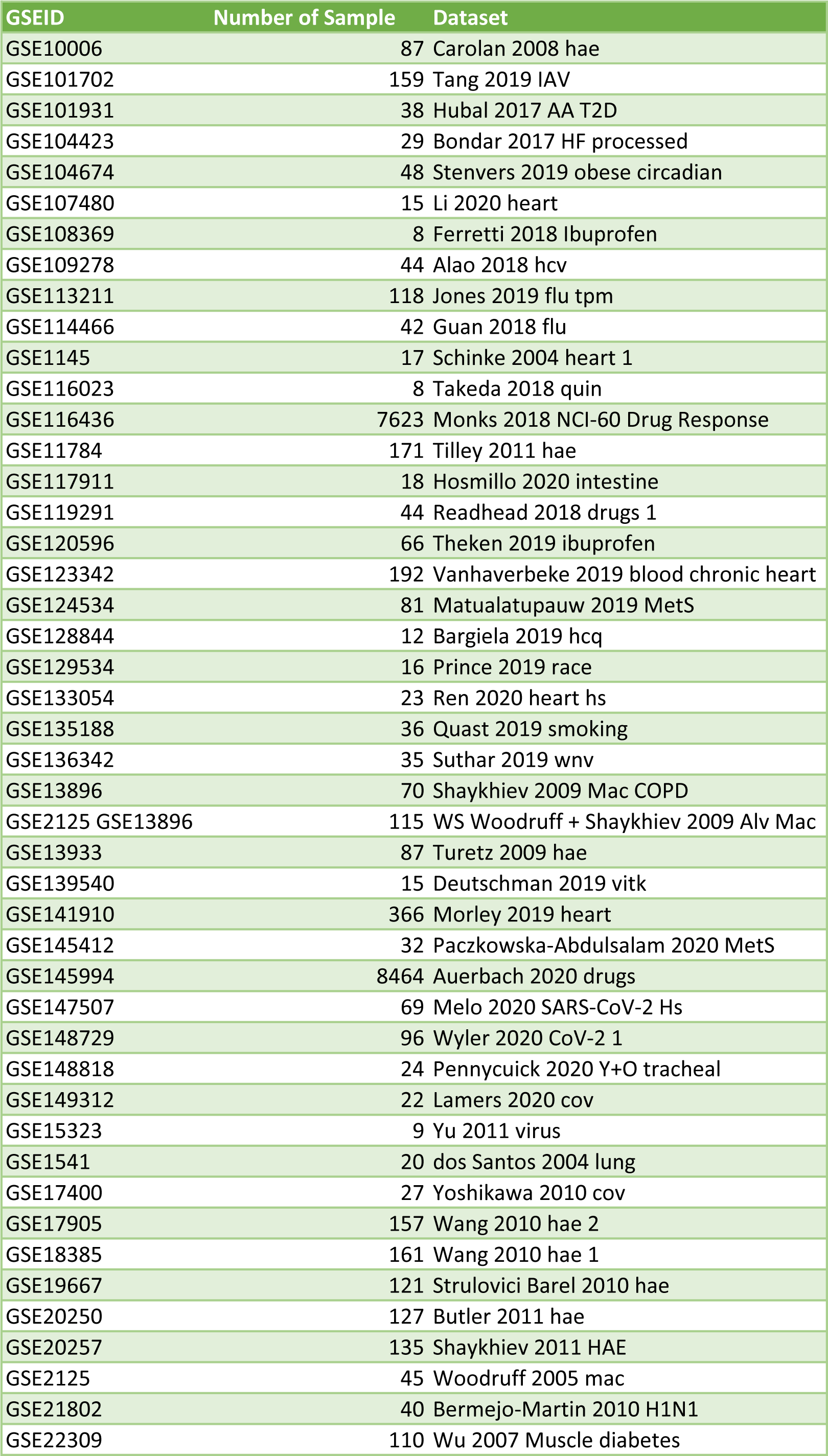
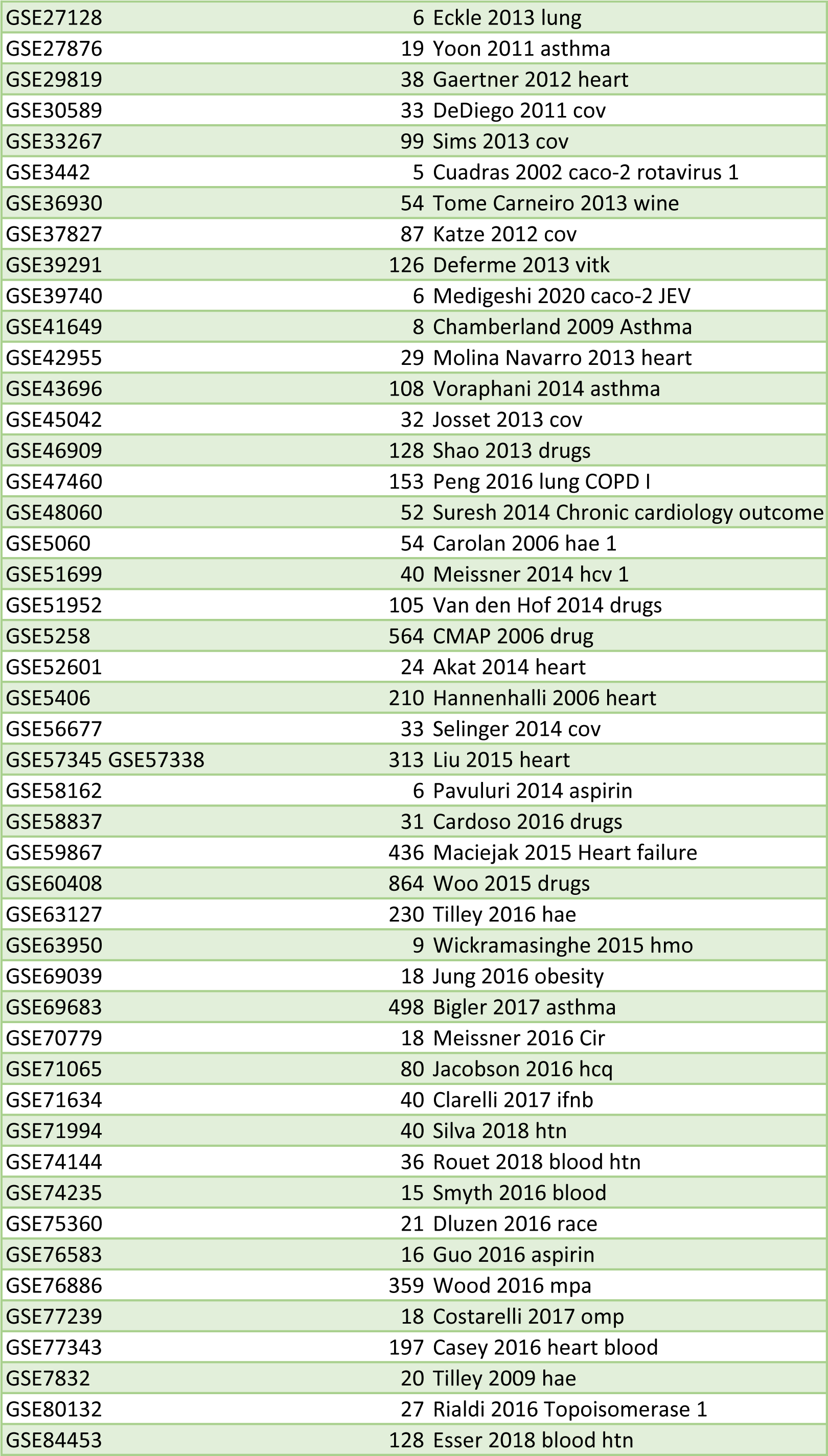
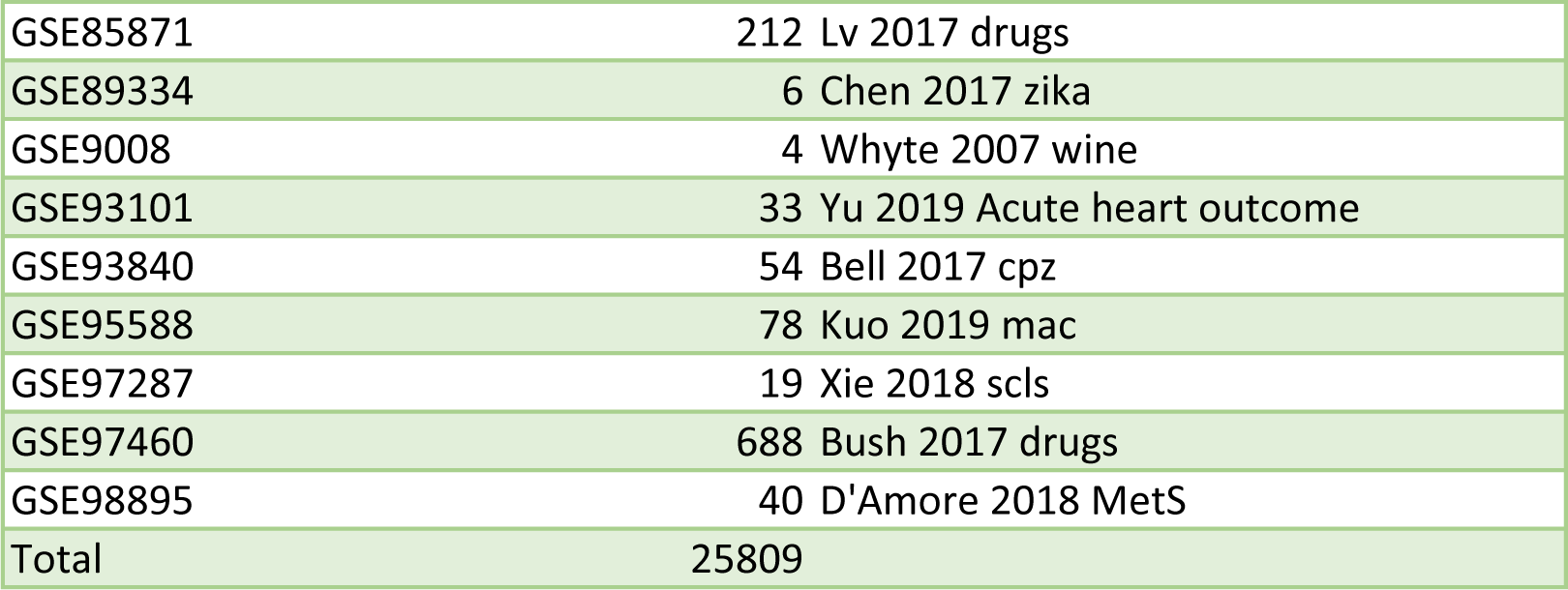
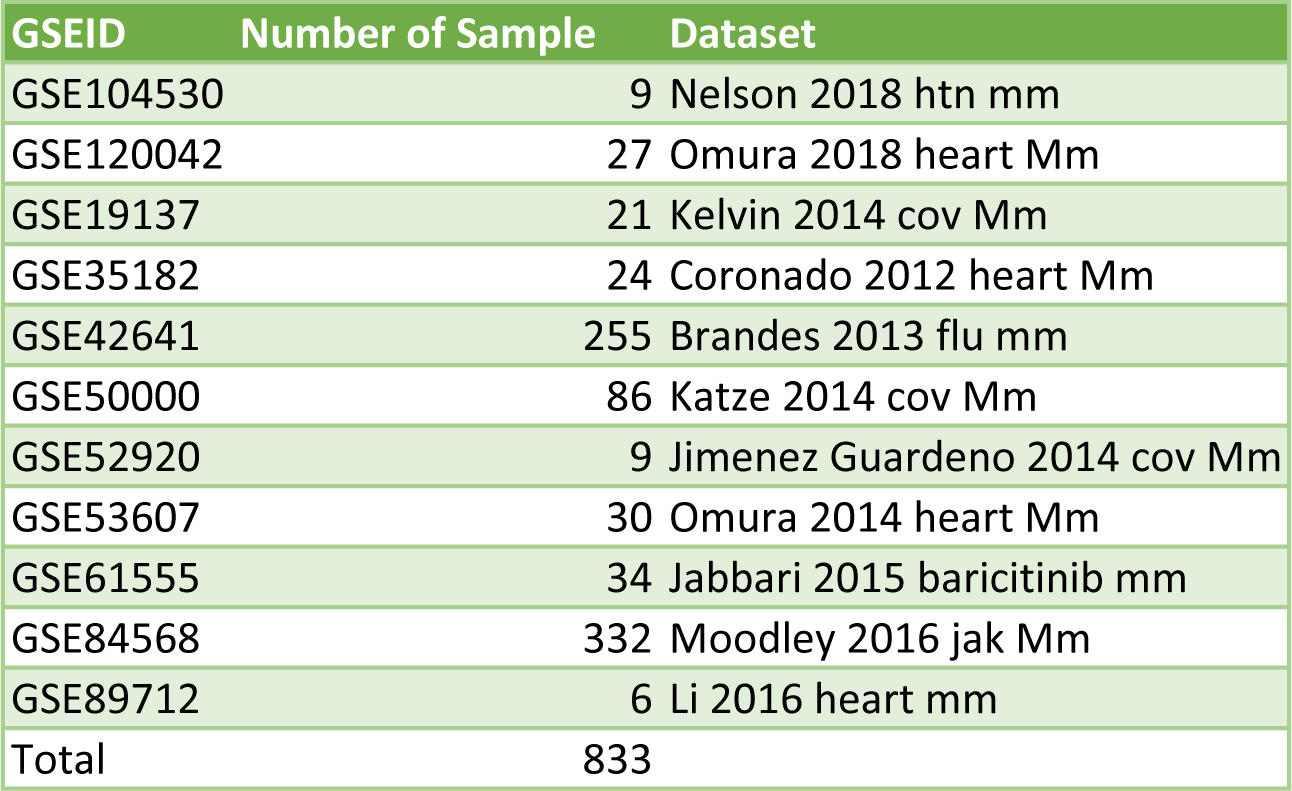
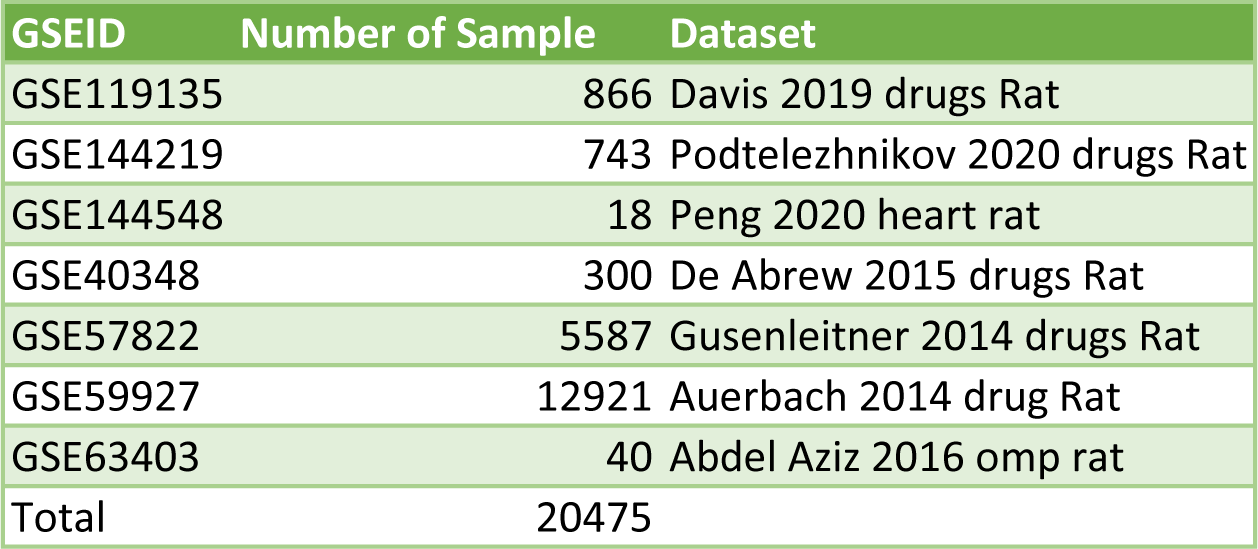

**Table S2.**
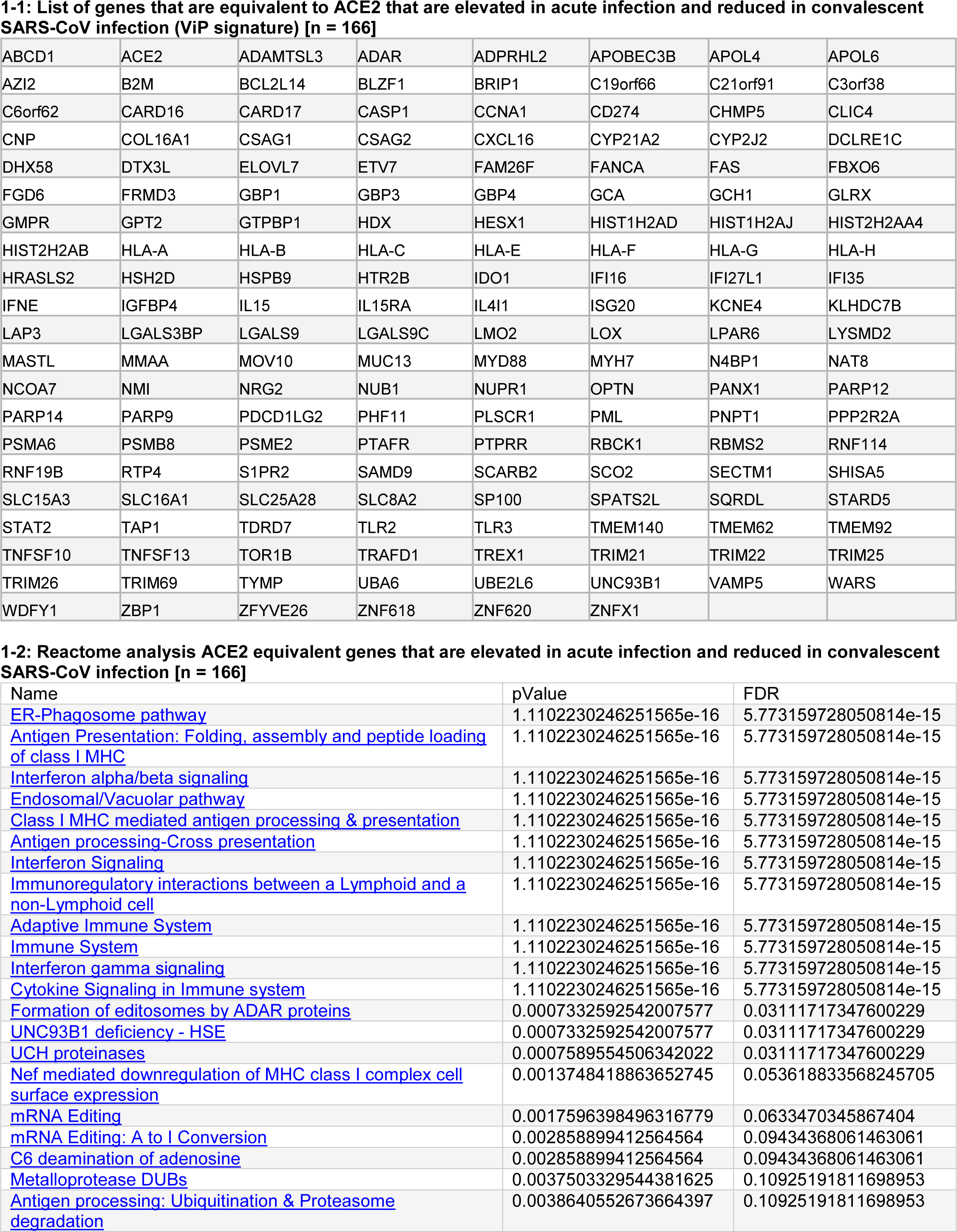
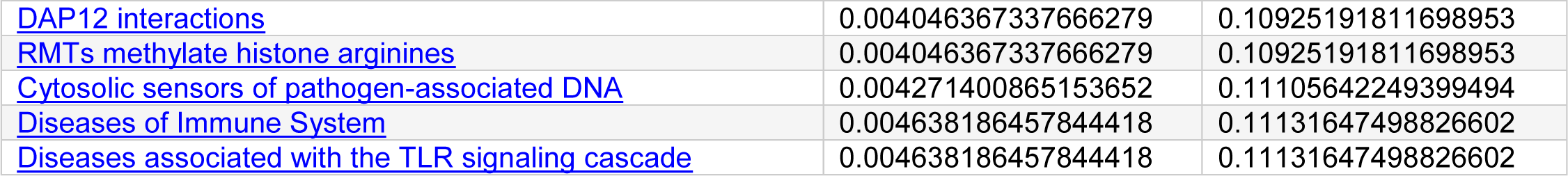
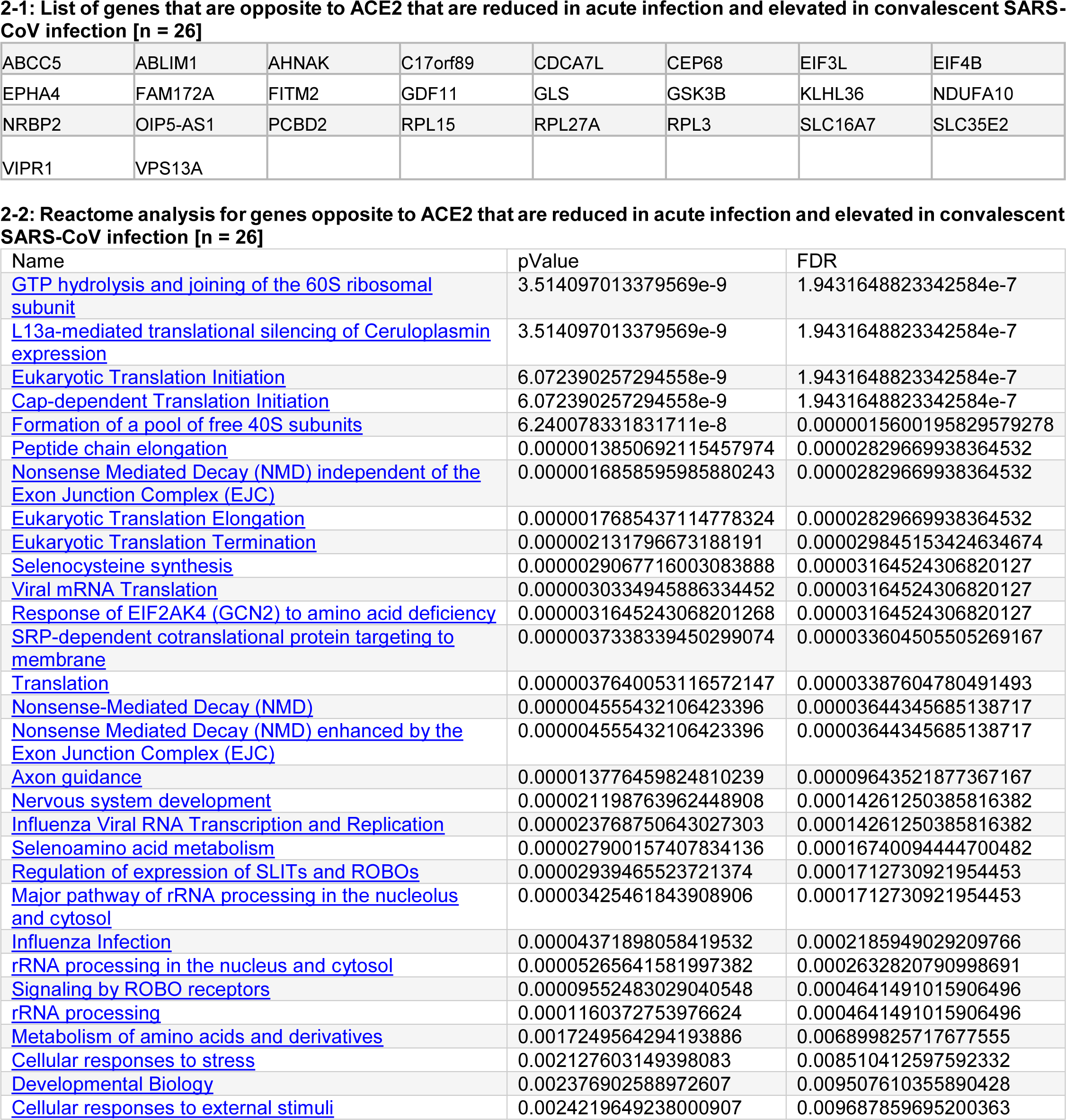
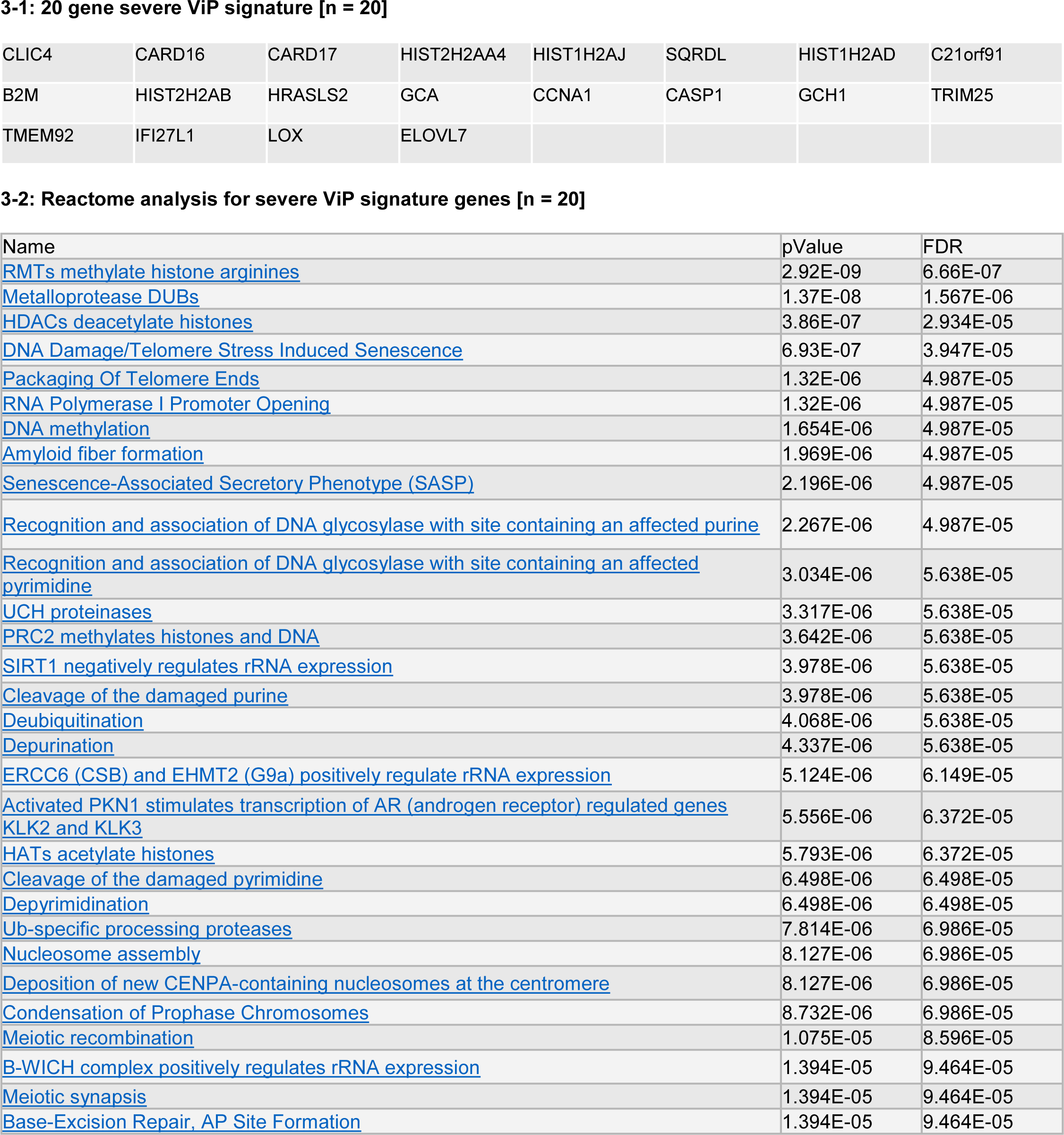

**Table S3.** Virus Infection Datasets Used in this Study.

| Virus | Type (RNA/DNA) | Portal of Entry (CME-dependent or independent) | GSE ID | In vitro/in vivo | Cell type | ROC AUC |
| --- | --- | --- | --- | --- | --- | --- |
| HIV | ss RNA (+) | CME-dependent | GSE125817 | in vitro | human monocyte derived dendritic cells (moDCs) | 1 |
| Zika | ss RNA (+) | CME-dependent | GSE146423 | in vitro | A549 cells (Adenocarcinomic human alveolar basal epithelial cells) | 1 |
| WNV | ssRNA (+) | CME-dependent | GSE138841 | in vitro | A549 cells (Adenocarcinomic human alveolar basal epithelial cells) | 1 |
| WNV | ssRNA (+) | CME-dependent | GSE136342 | in vitro | Human monocyte derived DCs moDCs | 1 |
| CoV (MERS) | ss RNA (+) | CME-dependent | GSE56677 | In vitro | Calu-3 2B4 cells; Human Coronavirus EMC 2012 (HCoV-EMC) | 1 |
| CoV (HCoV-EMC) | ss RNA (+) | CME-dependent | GSE45042 | In vitro | Calu-3 2B4 cells were infected with Human Coronavirus EMC 2012 (HCoV-EMC) | 1 |
| CoV | ss RNA (+) | CME-dependent | GSE17400 | In vitro | Calu3 cells with SARS-CoV1 and Dhori virus (DHOV), a member of the Orthomyxoviridae family within the Thogotovirus genus | 1 |
| CoV | ss RNA (+) | CME-dependent | GSE30589 | In vitro | Vero E6 cells SARS-CoV-1 | 1 |
| H1N1 | ss RNA (-) | CME-dependent | GSE47963 | In vitro | HAE cultures with H1N1 Inf virus | 1 |
| Inf A/B | ss RNA (-) | CME-dependent | GSE68310 | in vivo (human) | Peripheral blood cells | 0.97 |
| CoV (SARS-CoV-1) | ss RNA (+) | CME-dependent | GSE33267 | In vitro | Calu-3 cells were infected with either icSARS CoV or the icSARS deltaORF6 | 0.97 |
| EnV HHV-6 | ds DNA | CME-independent | GSE40396 | in vivo (human) | Peripheral blood cells | 0.95 |
| CoV (SARS-CoV-1) | ss RNA (+) | CME-dependent | GSE37827 | In vitro | Calu-3 cells were infected with either icSARS CoV or the cSARS Bat SRBD | 0.94 |
| HIV | ss RNA (+) | CME-dependent | GSE140713 | in vivo (human) | PBMCs | 0.93 |
| HCV | ss RNA (+) | CME-dependent | GSE40184 | in vivo (human) | PBMCs | 0.93 |
| HSV | ds DNA | CME-independent | GSE46042 | in vitro | Pluripotent stem cell derived neuronallineages | 0.89 |
| Ebola | ss RNA (-) | CME-dependent (and Macropinocytosis) | GSE130629 | in vivo (mouse) | Liver and spleen | 0.88 |
| HAV | ss RNA (+) | CME-dependent | GSE40396 | in vivo (human) | Peripheral blood cells | 0.87 |
| CMV | ds DNA | CME-independent | GSE81246 | in vivo (human) | PBMCs | 0.86 |
| SARS-CoV-2 | ss RNA (+) | CME-dependent | GSE147507 | In vitro | Primary human lung epithelium (NHBE) and transformed lung alveolar (A549) cells were mock treated or infected with SARS-CoV-2 (USA-WA1/2020) | 0.86 |
| HHV6 | ds DNA | CME-independent | GSE40396 | in vivo (human) | Peripheral blood cells | 0.85 |
| HEV | ss RNA (+) | CME-dependent | GSE36539 | in vivo (human) | Peripheral blood cells | 0.85 |
| VZV | ds DNA | CME-independent | GSE136586 | in vitro | Primary keratinocytes | 0.81 |
| SARS-CoV1 | ss RNA (+) | CME-dependent | GSE47963 | In vitro | HAE cultures with SARS-CoV1 | 0.81 |
| VZV | ds DNA | CME-independent | GSE54385 | in vitro | human dermal fibroblasts and neuron | 0.75 |
| HBV | ds DNA | CME-dependent | GSE135501 | in vivo (human) | Peripheral blood cells | 0.71 |
| AdV | ss DNA | CME-dependent | GSE40396 | in vivo (human) |  | 0.71 |
| HRV | ss RNA (+) | CME-dependent | GSE40396 | in vivo (human) |  | 0.7 |
| CMV | ds DNA | CME-independent | GSE99454 | in vitro | MRC-5 and ARPE-19 cells | 0.65 |
| EBV | ds DNA | CME-dependent | GSE135644 | in vitro | AGS and AGS-EBV cells | 0.64 |
| Measles | ss RNA (-) | CME-independent (Caveolin-dependent) | GSE5808 | in vivo | PBMC | 0.53 |
| HPV | ds DNA | CME-independent | GSE137965 | in vitro | Primary keratinocytes | 0.51 |

**Table S4.** Table of 20 genes that define “severity” within the 166-gene ViP signature.

| Gene Name | Host responses/infection | Targeted pathway | Drug/biologics-FDA approved |
| --- | --- | --- | --- |
| **CLIC4** | Chloride intracellular channels 4 is required for the replication of Chikungunya virus (PMID: 31483794) | - | Ribavirin PMID: 31483794 |
| **CARD16** | CARD 16 increases after the influenza A virus (PMID: 29299535) | Therapeutics on NLRP3<br>NLRP3 inhibitor— | Canakinumab<br>Anakinra<br>rilonacept |
| **CARD17** | CARD 17 increases after Chikungunya virus (CHIKV) infection (PMID: 31211814) & after the influenza A virus (PMID: 29299535) | Therapeutics on NLRP3<br>NLRP3 inhibitor— | Canakinumab<br>Anakinra<br>rilonacept |
| **HIST2H2AA4** | Histone family | pan-histone deacetylase (HDAC)<br>inhibitor SAHA | SAHA |
| **HIST1H2AJ** | Histone family | pan-histone deacetylase (HDAC)<br>inhibitor SAHA | SAHA |
| **SQRDL** | Sulfide quinone reductase |  |  |
| **HIST1H2AD** | Histone family | pan-histone deacetylase (HDAC)<br>inhibitor SAHA | SAHA |
| **C21orf91** | Cold Sore Susceptibility Gene 1, influences pathogenesis of herpes virus (PMID: 27081513) | - |  |
| **B2M** | b2 microglobulin mRNA levels are transiently increased after infection with Neurovirulent Sindbis Virus, suggesting that transcription of these mRNAs are coordinately regulated in neurons. (PMCID: PMC112110) |  |  |
| **HIST2H2AB** | Histone family | pan-histone deacetylase (HDAC)<br>inhibitor SAHA | SAHA |
| **HRASLS2** | Phospholipase A And Acyltransferase 2 |  |  |
| **GCA** | Grancalcin/GCA heterodimerization of TLR9 is important for TLR9-mediated downstream signaling during fine tune processes against viral infection (PMID: 26648480) |  |  |
| **CCNA1** | Cyclin A1 |  |  |
| **CASP1** | Caspase 1 |  |  |
| **GCH1** | GTP cyclohydrolase;<br>Neopterin is a biomarker for viral infection<br>It is produced as a by-product in tetrahydrobiopterin (BH4) de novo synthesis and mirrors the activity of the rate limiting enzyme in the BH4 synthesis cascade |  |  |
| **TRIM25** | tripartite motif containing 25 (TRIM25) is an E3 ubiquitin ligase and activates RIG1. It is involved in innate immune responses regulates intracellular signaling and/or RNA virus replication. PMID: 29018447 |  |  |
| **TMEM92** | Transmembrane Protein 92 |  |  |
| **IFI27L1** | Interferon Alpha Inducible Protein 27 Like 1 |  |  |
| **LOX** | Lipoxygenase increased in lung cells during RSV infection. It also gives higher CCL3 and CCL5 (PMID: 28579398) | Maraviroc prevent the function of CCR5 receptor and it will block CCL5 binding (PMID: 22637726) | Maraviroc |
| **ELOVL7** | host fatty acid elongase 7; required for efficient virus release and virion infectivity (PMID: 25732827) |  |  |

**Table S5.** Demographics of the UCSD COVID-19 cohort participants for plasma.

| UCSD ID | Acute/<br>Convalescent | Days post<br>symptom<br>onset | Age | Gender | Peak Disease Severity |
| --- | --- | --- | --- | --- | --- |
| CoV3 | Acute | 37 | 59 | M | Critical |
| CoV23 | Acute | 18 | 72 | F | Moderate |
| CoV36 | Acute | 17 | 43 | F | Critical |
| CoV37 | Acute | 4 | 25 | F | Mild |
| CoV38 | Acute | 4 | 32 | M | Mild |
| CoV41 | Acute | 7 | 83 | M | Mild |
| CoV40 | Acute | 10 | 83 | M | Severe to Critical (NRB in ICU but never intubated) |
| CoV42 | Acute | 6 | 58 | M | Mild to Moderate |
| CoV64 | Acute | 13 | 62 | M | Moderate |
| CoV65 | Acute | 28 | 42 | M | Critical |
| CoV66 | Acute | 6 | 53 | F | Moderate |
| Cov77 | Acute | 9 | 81 | M | Mod-Severe |
| Cov78 | Acute | 10 | 42 | M | Mod-Severe |
| CoV81 | Acute | 22 | 37 | F | Critical |
| CoV82 | Acute | 8 | 53 | M | Critical |
| CoV92 | Acute | 10 | 84 | M | Critical |
| CoV93 | Acute | 4 | 41 | M | Critical |
| CoV97 | Acute | 13 | 65 | M | Critical |
| CoV98 | Acute | 5 | 86 | M | Critical |
| CoV99 | Acute | 22 | 77 | F | Critical |
| CoV14 | Convalescent | 24 | 32 | M | Mild |
| CoV21 | Convalescent | 27 | 42 | M | Mild |
| CoV22 | Convalescent | 34 | 51 | M | Mild |
| CoV25 | Convalescent | 21 | 40 | F | Mild |
| CoV26 | Convalescent | 30 | 43 | F | Mild |
| CoV29 | Convalescent | 30 | 55 | F | Mild |
| CoV30 | Convalescent | 27 | 40 | M | Mild to Moderate |
| CoV31 | Convalescent | 33 | 64 | M | Mild |
| Cov7 | Convalescent | 25 | 44 | F | mild |
| Cov8 | Convalescent | 22 | 45 | M | mild |
| Cov9 | Convalescent | 25 | 43 | F | mild |
| Cov12 | Convalescent | 25 | 39 | F | mild |
| Cov18 | Convalescent | 23 | 48 | F | mild |
| Cov62 | Convalescent | 50 | 55 | M | mild |
| Cov63 | Convalescent | 39 | 68 | M | mild to moderate |
| Cov64 | Convalescent | 56 | 44 | F | mild |
| Cov201 | Acute | 12 | 73 | M | moderate |
| Cov204 | Acute | 13 | 54 | M | moderate |
| Cov205 | Acute | 9 | 37 | M | moderate |
| Cov206 | Acute | 10 | 77 | F | moderate |
| Cov207 | Acute | 8 | 49 | M | moderate |
| Cov210 | Acute | 5 | 48 | M | moderate |
| Cov211 | Acute | 8 | 39 | F | moderate |
| Cov202 | Acute | 10 | 37 | M | severe |
| Cov106 | Convalescent | 74 | 59 | M | Moderate |
| Cov103 | Convalescent | 79 | 50 | M | mild to moderate |
| Cov208A | Acute | 7 | 84 | M | Fatal |
| Cov213A | Acute | NA | 37 | M | Asymptomatic |
| Cov215 | Acute | 9 | 68 | M | Severe |
| Cov218 | Acute | 5 | 53 | M | Severe |
| CoV220 | Acute | N/A | 55 | M | Asymptomatic |
| CoV221 | Acute | 5 | 68 | M | moderate to severe |
| CoV222 | Acute | 5 | 68 | M | moderate |
| CoV223 | Acute | 7 | 57 | M | moderate to severe |

**Table S6.** Demographics of the UCSD COVID-19 cohort participants for lung tissue.

| Sample Name | Age | Gender | Days spent hospitalized COVID + until expired | Smoking history | Drugs administered during hospitalization | Subsequent infection? | Reason for surgery (NON SARS-CoV-2) | Histology (NON SARS-CoV-2) |
| --- | --- | --- | --- | --- | --- | --- | --- | --- |
| Autopsy 2 | 64 | Male | 35 days | Smoked 1.5-2 Packs per day (PPD) until 1988 when they quit. Smoked cannabis daily | Lasix (furosemide), Amiodarone HCL, stress dose steroids, vancomycin/zosyn, IV ceftriaxone, norepinephrine and vasopressin, digoxin, vecuronium | MSSA from being intubated |  |  |
| Autopsy 3 | NA | NA | NA | NA | NA | NA |  |  |
| Autopsy 4 | 90 | Female | 3 | NA | vancomycin/zosyn, low dose of norepinephrine | none |  |  |
| Autopsy 5 | 57 | Male | 28 | NA | Cefepime, Hydroxychloroquine, vancomycin, Norepinephrine, vasopressin, esmolol, Unasyn, bivalirudin, pressors (levo, vaso, phenylephrine), linezolid, Tocilizumab v. Placebo (information was not given what group he belonged to) | E.feacalis and klebsillea PNA from tracheal aspirate |  |  |
| JA 5 (NON SARS-CoV-2) | 46 | Female | NA | Non-smoker | NA | NA | Left Lower Lobe Nodule | Invasive Adenocarcinoma |
| JA 6 (NON SARS-CoV-2) | 70 | Male | NA | Non-smoker | NA | NA | Left Lower Lobe Nodule | Granulomatous-like inflammation and scar |

