## Supplementary Online Materials for "AI-guided discovery of the invariant host response to viral pandemics"

### Affiliations:

*Equal contribution
† Co-Corresponding

### Corresponding authors:

**SUPPLEMENTARY MATERIALS: Includes**

- Detailed Materials and Methods
- Supplementary Text- **n/a**
- Figures **S1-S4**
- Tables **S1-6**
- External Databases **- None**
- References ***(1-20)***

### DETAILED SUPPLEMENTARY METHODS:

#### Data Collection and Annotation

Publicly available microarray and RNASeq databases were downloaded from the National Center for Biotechnology Information (NCBI) Gene Expression Omnibus (GEO) website[1-3]. Gene expression summarization was performed by normalizing Affymetrix platforms by RMA (Robust Multichip Average)[4, 5] and RNASeq platforms by computing TPM (Transcripts Per Millions)[6, 7] values whenever normalized data were not available in GEO. We used log2(TPM) if TPM > 1 and (TPM – 1) if TPM < 1 as the final gene expression value for analyses. A catalog of all datasets analyzed in this work can be found in **Supplementary Table 1**.

#### Rapid autopsy procedure for tissue collection

The lung specimens from the COVID 19 positive human subjects were collected using autopsy (study was IRB Exempt). All donations to this trial were obtained after telephone consent followed by written email confirmation with next of kin/power of attorney per California state law (no in-person visitation could be allowed into our COVID-19 ICU during the pandemic).

The team member followed the CDC guidelines for COVID19 and the autopsy procedures[8, 9]. Lung specimens were collected in 10% Zinc-formalin and stored for 72 h before processing for histology. Patient characteristic is listed in **Supplementary Table 6**.

Autopsy 2 was a standard autopsy performed by anatomical pathology in the BSL3 autopsy suite. The patient expired and his family consented for autopsy. After 48 hours, lungs were removed and immersion fixed whole in 10% formalin for 48 hours and then processed further. Lungs were only partially fixed at this time (about 50% fixed in thicker segments) and were sectioned further into small 2-4cm chunks and immersed in 10% formalin for further investigation.

Autopsy 4 and 5 were collected from rapid postmortem lung biopsies. The procedure was performed in the Jacobs Medical Center ICU (all of the ICU rooms have a pressure-negative environment, with air exhausted through HEPA filters [Biosafety Level 3 (BSL3)] for isolation of SARS-CoV-2 virus). Biopsies were performed 2-4 hours after patient expiration. Ventilator was shut off to reduce aerosolization of viral particles at least 1 hour after loss of pulse and before the sample collection. Every team member had personal protective equipment in accordance with the University policies for procedures on patients with COVID-19 (N95 mask + surgical mask, hairnet, full face shield, surgical gowns, double surgical gloves, booties). Lung biopsies were obtained after L-thoracotomy in the 5th intercostal space by our cardiothoracic surgery team. Samples were taken from the left upper lobe (LUL) and left lower lobe (LLL) and then sectioned further.

**COVID-19 donors**

Blood from COVID-19 donors was either obtained at a UC San Diego Health clinic under the approved IRB protocols of the University of California, San Diego (UCSD; 200236X) or recruited at the La Jolla Institute under IRB approved (LJI; VD-214). COVID-19 donors were California residents, who were either referred to the study by a health care provider or self-referred. Blood was collected in acid citrate dextrose (ACD) tubes (UCSD) or in EDTA tubes (LJI) and stored at room temperature prior to processing for plasma collection. Seropositivity against SARS-CoV-2 was confirmed by ELISA. At the time of enrollment, all COVID-19 donors provided written informed consent to participate in the present and future studies. Patient characteristic is listed in **Supplementary Table 5**.

**Plasma isolation**

Whole blood was collected in heparin coated blood bags (healthy unexposed donors) or in ACD tubes (COVID-19 donors) and centrifuged for 25 min at 1850 rpm to separate the cellular fraction and plasma. The plasma was then carefully removed from the cell pellet and stored at -80C.

**Animal Study**

Lung samples from 8-week-old Syrian hamsters were generated from experiments conducted exactly as in a previously published study[10]. Animal studies were approved and performed in accordance with Scripps Research IACUC Protocol #20-0003 (PI: Tom Rogers, PMID: 32540903). We chose three different groups of samples: uninfected control, SARS-CoV-2 challenge after Den3 (antibody to dengue virus), and SARS-CoV-2 challenge after Anti-CoV2 (CC12.2; a potent SARS-CoV-2 neutralizing antibodies)[10].

**Plasma IL15 cytokine ELISA**

Plasma obtained from COVID-19 patients were used to quantify IL15 cytokine using ELISA MAX Deluxe set (Biolegend Cat. No. 435104) according to the manufacturer's recommended protocol.

The concentrations of IL15 cytokine were compared using Welch's t-test. A $p < 0.05$ denoted statistical significance.

**Multiplex measurement of human serum cytokines**

Human serum cytokines measurement was performed using customized Meso Scale Discovery (MSD)V-PLEX sandwich immunoassays. Human serum samples separated from peripheral blood of COVID-19 patients and healthy volunteers were analyzed using customized standard multiplex plates as per the manufacturer's instructions.

**Immunohistochemistry**

COVID-19 samples were inactivated by storing in 10% formalin for 2 days and then transferred to zinc-formalin solution for another 3 days. The deactivated tissues were transferred to 70% ethanol and cassettes were prepared for tissue sectioning. The slides containing hamster and human lung tissue sections were deparaffinized in xylene (Sigma-Aldrich Inc., MO, USA; catalog# 534056) and rehydrated in graded alcohols to water. For IL15RA antigen retrieval, slides were immersed in Tris-EDTA buffer (pH 9.0) and boiled for 10 minutes at 100°C. Slides were immersed in Tris-EDTA-Tween 20 buffer (pH 9.0) and pressure cooked for 3 minutes, for IL15 antigen retrieval. Endogenous peroxidase activity was blocked by incubation with 3% H2O2 for 10 minutes. To block non-specific protein binding 2.5% goat serum (Vector Laboratories, Burlingame, USA; catalog# S-1012) was added. Tissues were then incubated with rabbit IL15RA polyclonal antibody (1:200 dilution; proteintech®, Rosemont, IL, USA; catalog# 16744-1-AP) for 1.5 hour and mouse IL15 monoclonal antibody (1:10 dilution; Santa Cruz Biotechnology, Inc., Dallas, TX, USA; catalog# sc-8437) at room temperature in a humidified chamber and then rinsed with TBS or PBS 3x, 5 minutes each. Sections were incubated with goat anti-rabbit (Vector Laboratories, Burlingame, USA; catalog# MP-7401) and goat anti-mouse (Vector Laboratories, Burlingame, USA; catalog# MP-7402) secondary antibodies for 30 minutes at room temperature and then washed with TBS or PBS 3x, 5 minutes each; incubated with DAB (Vector Laboratories, Burlingame, USA; catalog# SK-4105), counterstained with hematoxylin (Sigma-Aldrich Inc., MO, USA; catalog# MHS1), dehydrated in graded alcohols, cleared in xylene, and cover slipped. Epithelial and stromal components of the lung tissue were identified by staining duplicate slides in parallel with hematoxylin and eosin (Sigma-Aldrich Inc., MO, USA; catalog# E4009) and visualizing by Leica DM1000 LED (Leica Microsystems, Germany).

**IHC Quantification**

IHC images were randomly sampled at different 300x300 pixel regions of interest (ROI). The ROIs were analyzed using IHC Profiler[11]. IHC Profiler uses a spectral deconvolution method of DAB/hematoxylin color spectra by using optimized optical density vectors of the color deconvolution plugin for proper separation of the DAB color spectra. The histogram of the DAB intensity was divided into 4 zones: high positive (0 to 60), positive (61 to 120), low positive (121 to 180) and negative (181 to 235). High positive, positive, and low positive percentages were combined to compute the final percentage positive for each ROI. The range of values for the percent positive is compared among different experimental groups. IL15 staining showed too many ROIs with low final percent positive score. We subtracted these background noise by focusing on only ROIs with greater than 20% positive percentages.

**RNA sequencing**

RNA sequencing libraries were generated using the Illumina TruSeq Stranded Total RNA Library Prep Gold with TruSeq Unique Dual Indexes (Illumina, San Diego, CA). Samples were processed following manufacturer's instructions, except modifying RNA shear time to five minutes. Resulting libraries were multiplexed and sequenced with 100 basepair (bp) Paired End (PE100) to a depth of approximately 25-40 million reads per sample on an Illumina NovaSeq 6000 by the Institute of Genomic Medicine (IGM) at the University of California San Diego. Samples were demuxltiplexed using bcl2fastq v2.20 Conversion Software (Illumina, San Diego, CA). RNASeq data was processed using kallisto (version 0.45.0), Mesocricetus auratus genome (MesAur1.0) and human genome GRCh38 Ensembl version 94 annotation (Homo_sapiens GRCh38.94 chr_patch_hapl_scaff.gtf). Gene-level TPM values and gene annotations were computed using tximport and biomaRt R package. A custom python script was used to organize the data and log reduced using log2(TPM) if TPM > 1 and TPM - 1 if TPM <= 1. For the hamster study kallisto index was prepared on Mesocricetus_auratus.MesAur1.0.ncrna.fa.gz + Mesocricetus_auratus MesAur1.0 cdna.all.fa.gz. The raw data and processed data are deposited in Gene Expression Omnibus under accession no GSE157058 (Hamster) and GSE157059 (Ileum).

**StepMiner Analysis**

StepMiner is a computational tool that identifies step-wise transitions in a time-series data.[12] StepMiner performs an adaptive regression scheme to identify the best possible step up or down based on sum-of-square errors. The steps are placed between time points at the sharpest change between low expression and high expression levels, which gives insight into the timing of the gene expression-switching event. To fit a step function, the algorithm evaluates all possible step positions, and for each position, it computes the average of the values on both sides of the step for the constant segments. An adaptive regression scheme is used that chooses the step positions that minimize the square error with the fitted data. Finally, a regression test statistic is computed as follows:

$$F\ stat = \frac{\sum_{i=1}^{n}(\widehat{X_i} - \bar{X})^2/(m-1)}{\sum_{i=1}^{n}(X_i - \widehat{X_i})^2/(n-m)}$$

Where $X_i$ for $i = 1$ to $n$ are the values, $\widehat{X_i}$ for $i = 1$ to $n$ are fitted values. m is the degrees of freedom used for the adaptive regression analysis. $\bar{X}$ is the average of all the values: $\bar{X} = \frac{1}{n} * \sum_{j=1}^{n} X_j$. For a step position at k, the fitted values $\widehat{X_i}$ are computed by using $\frac{1}{k} * \sum_{j=1}^{n} X_j$ for $i = 1$ to $k$ and $\frac{1}{(n-k)} * \sum_{j=k+1}^{n} X_j$ for $i = k + 1$ to $n$.

**Boolean Analysis**

**Boolean logic** is a simple mathematic relationship of two values, i.e., high/low, 1/0, or positive/negative. The Boolean analysis of gene expression data requires the conversion of expression levels into two possible values. The ***StepMiner*** algorithm is reused to perform Boolean analysis of gene expression data.[13] **The Boolean analysis** is a statistical approach which creates binary logical inferences that explain the relationships between phenomena. Boolean analysis is performed to determine the relationship between the expression levels of pairs of genes. The ***StepMiner*** algorithm is applied to gene expression levels to convert them into Boolean values (high and low). In this algorithm, first the expression values are sorted from low to high and a rising step function is fitted to the series to identify the threshold. Middle of the step is used as the StepMiner threshold. This threshold is used to convert gene expression values into Boolean values. A noise margin of 2-fold change is applied around the threshold to determine intermediate values, and these values are ignored during Boolean analysis. In a scatter plot, there are four possible quadrants based on Boolean values: (low, low), (low, high), (high, low), (high, high). A Boolean implication relationship is observed if any one of the four possible quadrants or two diagonally opposite quadrants are sparsely populated. Based on this rule, there are six kinds of Boolean implication relationships. Two of them are symmetric: equivalent (corresponding to the positively correlated genes), opposite (corresponding to the highly negatively correlated genes). Four of the Boolean relationships are asymmetric and each corresponds to one sparse quadrant: (low => low), (high => low), (low => high), (high => high). BooleanNet statistics (**Figure 2A**) is used to assess the sparsity of a quadrant and the significance of the Boolean implication relationships[13, 14]. Given a pair of genes A and B, four quadrants are identified by using the StepMiner thresholds on A and B by ignoring the Intermediate values defined by the noise margin of 2 fold change (+/- 0.5 around StepMiner threshold). Number of samples in each quadrant are defined as $a_{00}$, $a_{01}$, $a_{10}$, and $a_{11}$ (Figure 1A) which is different from X in the previous equation of F stat. Total number of samples where gene expression values for A and B are low is computed using the following equations.

$$nA_{low} = (a_{00} + a_{01}), nB_{low} = (a_{00} + a_{10}),$$

Total number of samples considered is computed using following equation.

$$total = a_{00} + a_{01} + a_{10} + a_{11}$$

Expected number of samples in each quadrant is computed by assuming independence between A and B. For example, expected number of samples in the bottom left quadrant $e_{00} = \hat{n}$ is computed as probability of A low (($a_{00}$ + $a_{01}$)/total) multiplied by probability of B low (($a_{00}$ + $a_{10}$)/total) multiplied by total number of samples. Following equation is used to compute the expected number of samples.

$$n = a_{ij}, \hat{n} = (nA_{low}/total * nB_{low}/total) * total$$

To check whether a quadrant is sparse, a statistical test for ($e_{00} > a_{00}$) or ($\hat{n} > n$) is performed by computing $S_{00}$ and $p_{00}$ using following equations. A quadrant is considered sparse if $S_{00}$ is high ($\hat{n} > n$) and $p_{00}$ is small.

$$S_{ij} = \frac{\hat{n} - n}{\sqrt{\hat{n}}}$$

$$p_{00} = \frac{1}{2}\left(\frac{a_{00}}{(a_{00} + a_{01})} + \frac{a_{00}}{(a_{00} + a_{10})}\right)$$

A suitable threshold is chosen for $S_{00}$ > sThr and $p_{00}$ < pThr to check sparse quadrant. A Boolean implication relationship is identified when a sparse quadrant is discovered using following equation.

***Boolean Implication*** = ($S_{ij}$ > sThr, $p_{ij}$ < pThr)

A relationship is called Boolean equivalent if top-left and bottom-right quadrants are sparse.

*Equivalent* = $(S_{01} > sThr,\ P_{01} < pThr, S_{10} > sThr, P_{10} < pThr)$

Boolean opposite relationships have sparse top-right ($a_{11}$) and bottom-left ($a_{00}$) quadrants.

*Opposite* = $(S_{00} > sThr,\ P_{00} < pThr, S_{11} > sThr, P_{11} < pThr)$

Boolean equivalent and opposite are symmetric relationship because the relationship from A to B is same as from B to A. Asymmetric relationship forms when there is only one quadrant sparse (A low => B low: top-left; A low => B high: bottom-left; A high=> B high: bottom-right; A high => B low: top-right). These relationships are asymmetric because the relationship from A to B is different from B to A. For example, A low => B low and B low => A low are two different relationships.

A low => B high is discovered if the bottom-left ($a_{00}$) quadrant is sparse and this relationship satisfies following conditions.

*A low => B high* = $(S_{00} > sThr,\ P_{00} < pThr)$

Similarly, A low => B low is identified if the top-left ($a_{01}$) quadrant is sparse.

*A low => B low* = $(S_{01} > sThr,\ P_{01} < pThr)$

A high => B high Boolean implication is established if the bottom-right ($a_{10}$) quadrant is sparse as described below.

*A high => B high* = $(S_{10} > sThr,\ P_{10} < pThr)$

Boolean implication A high => B low is found if the top-right ($a_{11}$) quadrant is sparse using following equation.

*A high => B low* = $(S_{11} > sThr,\ P_{11} < pThr)$

For each quadrant a statistic $S_{ij}$ and an error rate $p_{ij}$ is computed. $S_{ij} > sThr$ and $p_{ij} < pThr$ are the thresholds used on the BooleanNet statistics to identify Boolean implication relationships.

Boolean analyses in the test dataset GSE47963 uses a threshold of sThr = 5 and pThr = 0.05. Boolean analysis on the large normal lung dataset GSE23546 uses a threshold of sThr = 6 and pThr = 0.1. These thresholds are more stringent compared to previously used thresholds sThr = 3 and pThr = 0.1 for BooleanNet[13, 15, 16] to focus on the strong candidates.

**BECC (Boolean Equivalent Correlated Clusters) Analysis**

BECC analysis[16] is based on Boolean Equivalent relationships, pair-wise correlation and linear regression analysis. BECC analysis begins with a seed gene. We used ACE2 as a seed gene in this paper. BECC analysis identified a set of genes Boolean Equivalent to ACE2 and Boolean Opposite to ACE2 using the BooleanNet statistic described above.

The BECC algorithm identified 367 genes 'Boolean Equivalent' and 163 genes 'Boolean Opposite' to the ACE2 gene. Reactome pathway analyses on both clusters showed that the 367-gene ACE2-equivalent cluster was enriched in viral response pathways and processes, whereas the 163-gene ACE2-opposite cluster represented housekeeping processes, implying that ACE2 and its related genes are drivers of host response in the setting of viral infections. These clusters were subsequently filtered using differential analysis on another dataset [GSE113211 (n = 118); **Fig 2B**] that profiled heterogeneous immunophenotypes of children with viral bronchiolitis (confirmed positive for the virus in ~100% patients; of which 25 % were infected with Influenza/Para-Influenza and 14.8% with human CoV). We chose GSE113211 (n=118) dataset to filter ViP genes because this is the only high-quality large in vivo dataset available with clinical annotation on two different tissue types: nasal mucosal scrapings (NMS) and PBMC. Transcriptomes were analyzed in nasal mucosal scrapings (NMS) and PBMC samples taken during an acute visit (AV) and during a subsequent visit at convalescence (CV)[17]. Of the 367 ACE2-equivalent genes, 166 genes (**Table S2**; 1-1) retained the "Boolean Equivalent" relationship with ACE2 *and* their expression was downregulated during the convalescence visit. Of the 163 ACE2-opposite genes, 26 genes (**Table S2;** 2-1) retained "Boolean Opposite" relationships with ACE2 *and* their expression were upregulated during the convalescence visit. All subsequent analyses were performed using the 166

–gene signature that had Boolean Equivalent relationship with ACE2 and that was down-regulated during a convalescent visit after acute viral bronchiolitis.

A gene signature score is computed using the 166-genes that were equivalent to ACE2 which is used to order the sample. To compute the ViP signature, first the genes present in this list were normalized according to a modified Z-score approach centered around StepMiner threshold (formula = (expr -SThr)/3*stddev). The normalized expression values for every probeset for 166 genes were added together to create the final ViP signature. The samples were ordered based on the final ViP signature. To compute the severe ViP signature, 166 genes were first ordered using T test between the mild vs severe cases in GSE101702 dataset, and top 20 genes (**Table S2;** 3-1) were selected from this list. We chose GSE101702 dataset to select Severe ViP genes because this is the only high-quality large dataset available with clinical severity annotation.

To test the significance of the ViP and Severe ViP genes, we subsampled GSE47963 dataset to see if similar number of genes appear after BECC analysis. We selected 250 samples from 438 total number of samples 10,000 times randomly and performed BECC on them. Boolean analysis on the original GSE47963 used thresholds of sThr = 5 and pThr = 0.05. These thresholds need to be adjusted when the number of samples are reduced. For the BECC analysis on the 250 randomly selected samples we used thresholds of sThr = 4 and pThr = 0.06 which discovers around 367 genes on average. Analysis of the genes discovered in the 250 subsampled datasets revealed that on average 87% (321 out 367) emerge again and 41 new genes appear. In this test, on average 90% (150 out of 166) ViP genes and 90% (18 out of 20) Sever ViP genes emerge again. 17 new ViP genes and 2 new Severe ViP genes appear. These results suggests that there are about 10% variation in the genes which is a reasonable criterion for robustness.

**Single Cell RNASeq data analysis**

Single Cell RNASeq data from GSE145926 and GSE150728 was downloaded from Gene Expression Omnibus (GEO) in the HDF5 Feature Barcode Matrix Format. The filtered barcode data matrix was processed using Seurat v3 R package[18]. B cells (CD19, MS4A1, CD79A), T cells (CD3D, CD3E, CD3G), CD4 T cells (CCR7, CD4, IL7R, FOXP3, IL2RA), CD8 T cells (CD8A, CD8B), Natural killer cells (KLRF1), Macs Monos DCs (TYROBP, FCER1G), Epithelial (SFTPA1, SFTPB, AGER, AQP4, SFTPC, SCGB3A2, KRT5, CYP2F1, CCDC153, TPPP3) cells were identified using relevant gene markers using SCINA algorithm[19]. Pseudo bulk datasets were prepared by adding counts from the different cell subtypes and normalized using log(CPM+1).

#### AI guided discovery of invariant host response

BECC requires the depth of Boolean equivalent relationship as a parameter. For example, if ACE2 is Equivalent to X and X is Equivalent to Y but ACE2 is not necessarily equivalent to Y, depth of X is 1 and Y is 2. The depth controls how much the list of genes that are Boolean equivalent to ACE2 is expanded. This list of genes is converted to a gene expression score based on average of the normalized gene expression values as mentioned before. The strength of classification of uninfected and infected samples using this score is computed by the ROC-AUC measurement. We performed a regression to identify the best depth that predicts uninfected vs infected samples in the cohort GSE47963 (n = 438). We tested how the gene expression score distinguish uninfected and infected samples as they are annotated in many other independent datasets. Our confidence on the host response being invariant depend on having this test pass in all properly annotated cohorts without exceptions.

#### Survival Outcome in COVID-19

Hospital-free days analysis (45 days followup) of COVID-19 patients (GSE157103) limited to less than 70 years old using sViP signature (low and high group) is analyzed using Kaplan-Meier and Cox-proportional hazard approach. The threshold to separate high and low group was computed using StepMiner determined threshold + a noise margin. The noise margin for sViP signature was computed by computing the total dynamic range (max – min) divided by 65 to bring it to comparable levels of two-fold change noise margin seen in gene expression datasets. For the IL15 transcript analysis samples were limited to only males with less than 70 years old. IL15 transcripts were divided into high, intermediate and low levels by using StepMiner threshold +/- noise margin 1 which is two-fold change in log scale. Low levels of IL15 were associated with unusually adverse outcome. High and intermediate levels were compared to demonstrate the significance of IL15 in the context of our manuscript.

### Statistical Analyses

Gene signature is used to classify sample categories and the performance of the multi-class classification is measured by ROC-AUC (Receiver Operating Characteristics Area Under The Curve) values. A color-coded bar plot is combined with a density plot to visualize the gene signature-based classification. All statistical tests were performed using R version 3.2.3 (2015-12-10). Standard t-tests were performed using python scipy.stats.ttest_ind package (version 0.19.0) with Welch's Two Sample t-test (unpaired, unequal variance (equal_var=False), and unequal sample size) parameters. Multiple hypothesis correction were performed by adjusting *p* values with statsmodels.stats.multitest.multipletests (fdr_bh: Benjamini/Hochberg principles). The results were independently validated with R statistical software (R version 3.6.1; 2019-07-05). Pathway analysis of gene lists were carried out via the Reactome database and algorithm[20]. Reactome identifies signaling and metabolic molecules and organizes their relations into biological pathways and processes. Kaplan-Meier analysis is performed using lifelines python package version 0.14.6. Violin, Swarm and Bubble plots are created using python seaborn package version 0.10.1.

### SUPPLEMENTARY FIGURES AND LEGENDS

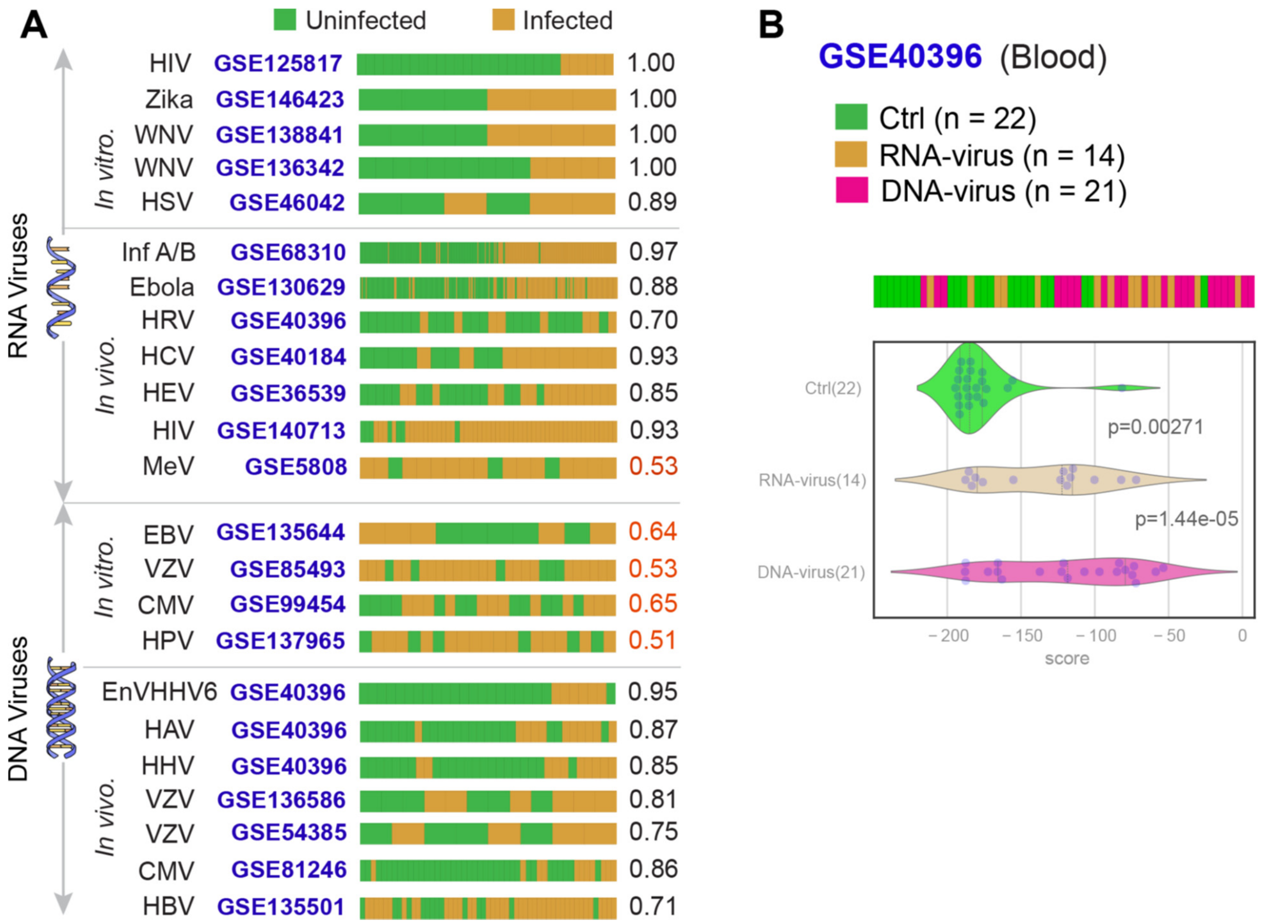


**Figure S1. The ViP signature distinguishes infected from uninfected samples across multiple RNA and DNA viruses. (A)** Bar plots showing the accuracy of the 166 gene ViP signature to classify infected vs. uninfected samples. Samples are categorized based on the nature of genetic material in the viruses (RNA *vs*. DNA) and whether the dataset was generated by assessing infections in *in vitro* cell-based models or in infected humans (*in vivo*). ROC-AUC values of infected samples classifications are shown on the right side of each bar plot. See also **Table S3**, which classifies these viruses based on their route of entry into cells, i.e., clathrin-dependent *vs*. independent endocytosis. ROC-AUC values of infected samples classifications are shown on the right side of each bar plot. **(B)** Bar (top) and violin (bottom) plots show that the ViP signature is equally effective in distinguishing RNA (R) and DNA (D) virus infections from uninfected controls (C) *in vivo*.

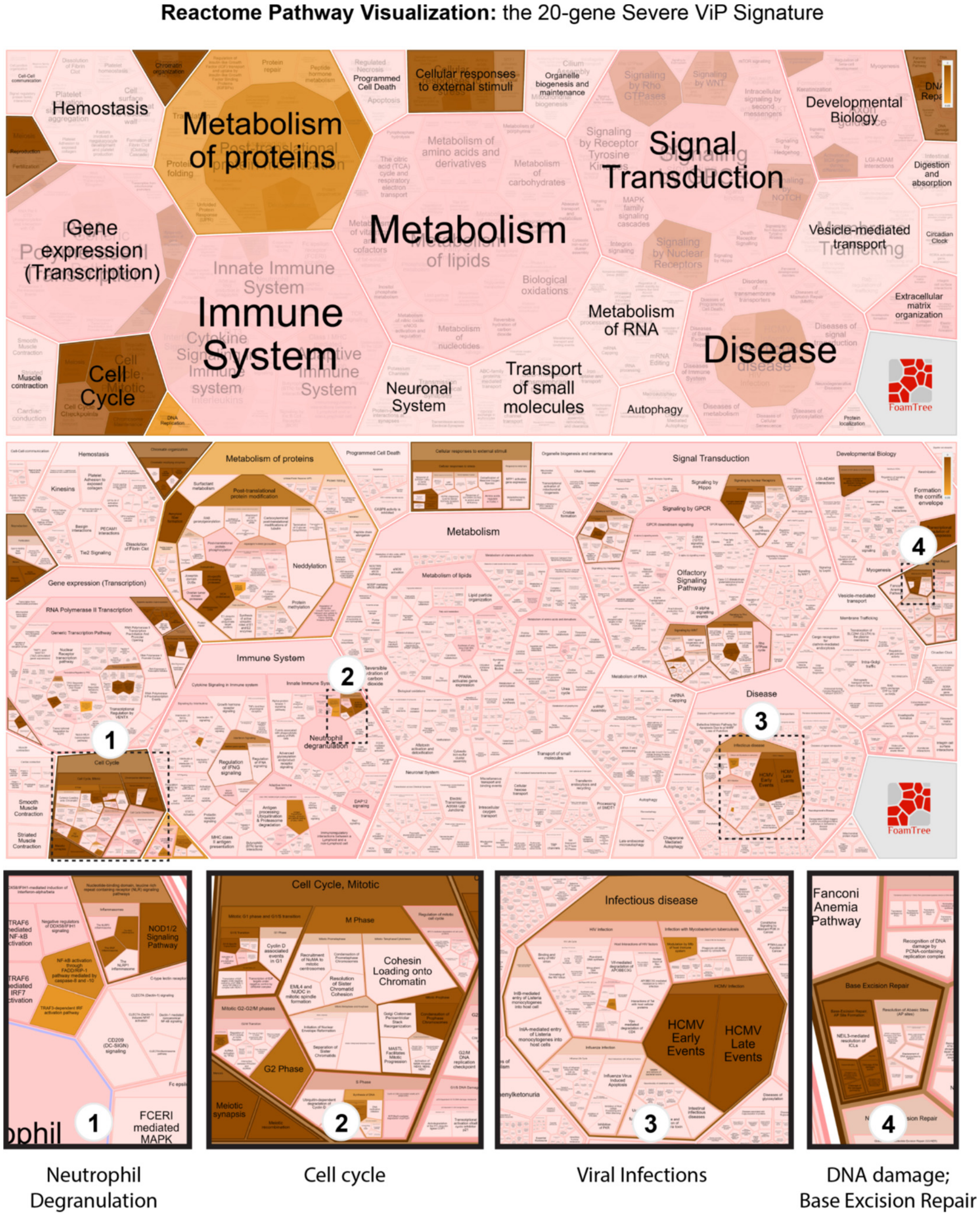


**Figure S2. Reacfoam analysis of 20-gene severe ViP signature. (A)** ReacFome pathway analysis of 20 gene severe ViP signature and visualization based on Voronoi tessellation. Cell cycle, Immune System, Disease and DNA Damage components are highlighted.

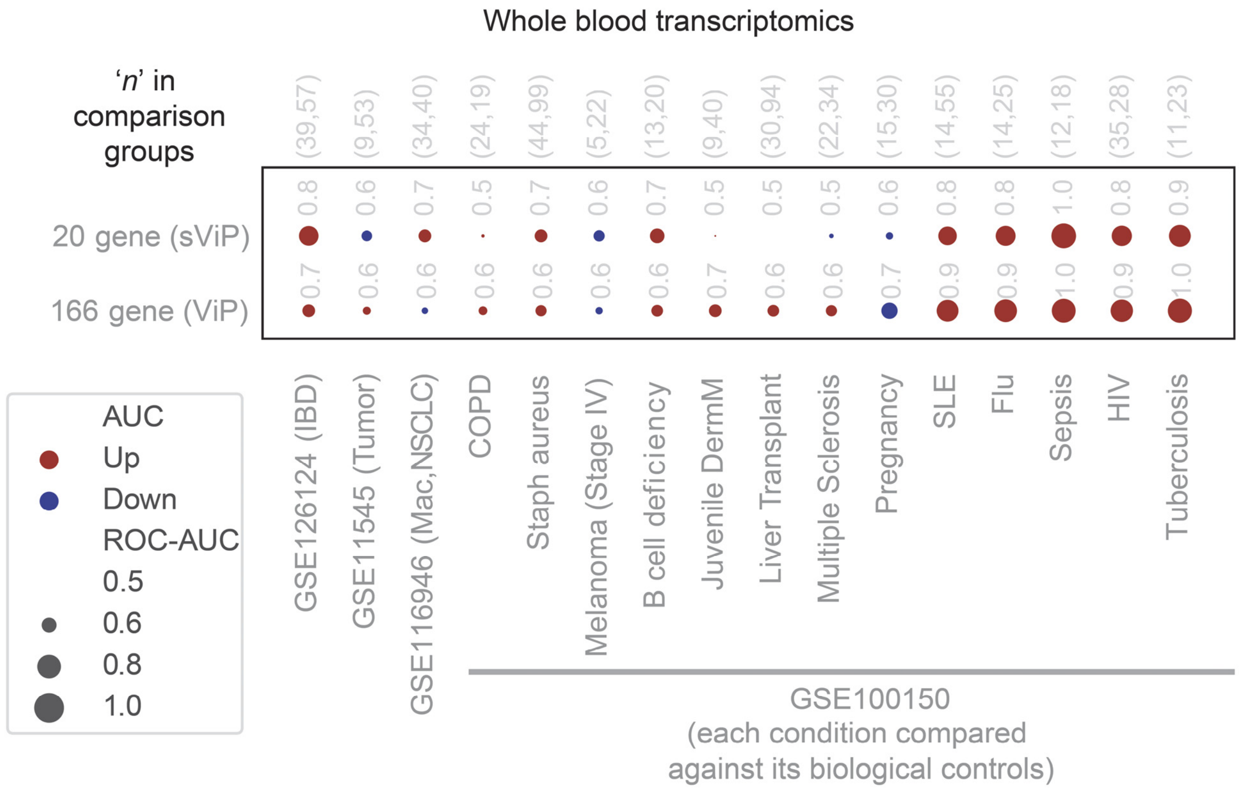


**Figure S3. ViP signatures are specific for diseases of infectious and inflammatory conditions.** Bubble plots showing up- (red) or downregulation (blue) of *ViP* signatures in blood samples from patients with diverse conditions and their corresponding controls. The size of bubble indicates the accuracy of classification (AUC ROC; see key) of controls from diseases samples using the signatures. Each dataset contains its own biological controls, and there are no replicates. IBD, Inflammatory bowel disease; NSCLC, non-small cell lung cancer; COPD, chronic obstructive pulmonary disease; Staph, staphylococcal infection; SLE, systemic lupus erythematosus.

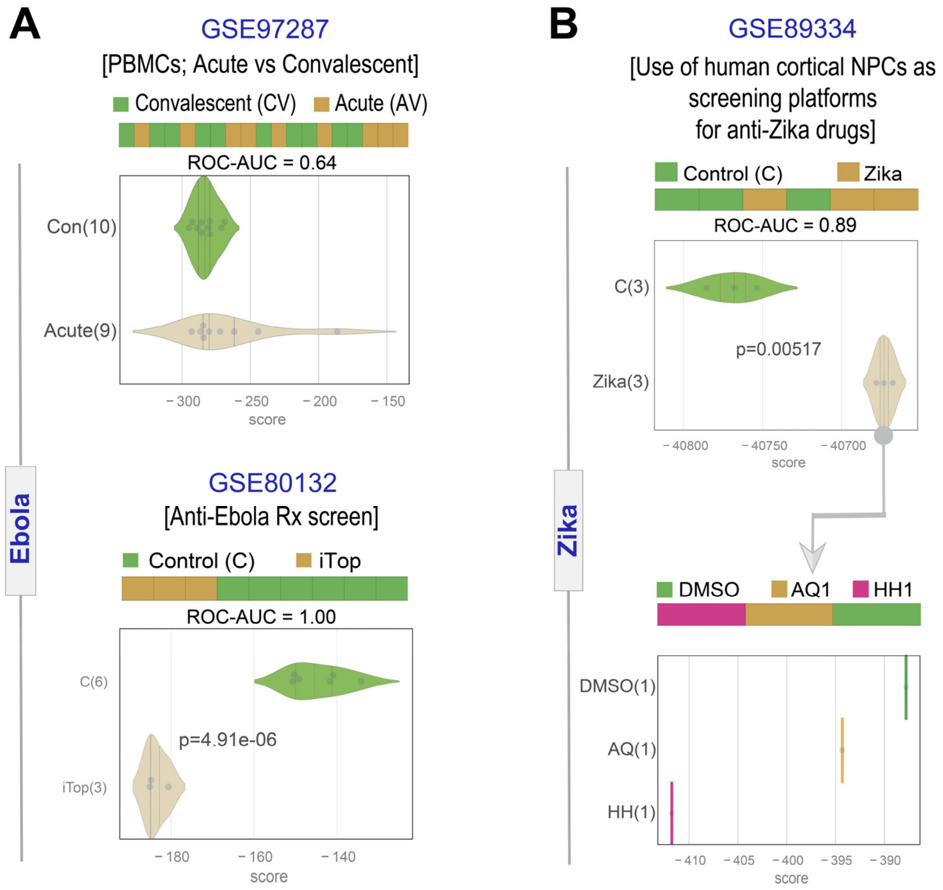


**Figure S4. Validation of ViP signature-guided therapeutic goals.**
**(A)** *Top*: 166-gene ViP signature-based classification of crisis and convalescence in PBMCs from patients with Ebola infection. *Bottom*: The effect of inhibiting Topoisomerase 1 (iTop) in a cultured cell line model infected in vitro with Ebola for the development of anti-Ebola therapeutics. **(B)** 166-gene ViP signature-based classification of human cortical neural progenitor cells infected *in vitro* with Zika virus (*top*) and the infected cells treated with two investigational drugs (*bottom*; two treatments, AQ1 and HH1) during screening assays.

### SUPPLEMENTARY TABLES INDEX: (Uploaded separately)

**Table S1:** Catalog of publicly available datasets analyzed in this work

**Table S2**: Gene clusters that constitute the ViP signature and their reactome analyses

**Table S3**: Classification of host response to viral infection using the ViP signature

**Table S4**: Table of 20 genes that define "severity" within the 166-gene ViP signature

**Table S5**. Demographics of the UCSD COVID-19 cohort participants for plasma.

**Table S6**. Demographics of the UCSD COVID-19 cohort participants for lung tissue.

11. Varghese F, Bukhari AB, Malhotra R, De A. IHC Profiler: an open source plugin for the quantitative evaluation and automated scoring of immunohistochemistry images of human tissue

samples. PLoS One. 2014;9(5):e96801. Epub 2014/05/08. doi: 10.1371/journal.pone.0096801. PubMed PMID: 24802416; PMCID: PMC4011881.
